# The molecular anatomy of mouse skin during hair growth and rest

**DOI:** 10.1101/750042

**Authors:** Simon Joost, Karl Annusver, Tina Jacob, Xiaoyan Sun, Unnikrishnan Sivan, Tim Dalessandri, Inês Sequeira, Rickard Sandberg, Maria Kasper

## Abstract

Skin homeostasis is orchestrated by dozens of cell types that together direct stem cell renewal, lineage commitment and differentiation. However, a systematic molecular atlas of full-thickness skin is lacking. Here, we used single-cell RNA-sequencing and mRNA-FISH to determine gene-expression identity and spatial location of skin cells during hair growth and rest. We defined 55 cell populations and made striking discoveries about the outer root sheath (ORS) and inner hair follicle layers that together coordinate hair production. The ORS is composed of two distinct cell types, companion layer cells resemble ORS and not inner layer cells, and we identified an asymmetric inner-layer structure with ORS cell identity. Moreover, the inner layers branch from transcriptionally uncommitted progenitors, and each lineage differentiation passes through an intermediate state. Altogether, we generated a comprehensive atlas with molecular and spatial information on epithelial and stromal cells, including fibroblasts, vascular and immune cells, that will spur new discoveries in skin biology.

**HIGHLIGHTS:**

- Comprehensive single-cell transcriptome atlas of full-thickness skin
- Outer root sheath (ORS) is composed of two distinct cell types
- Companion layer transcriptionally resembles ORS
- Transcriptional reconstruction of the internal hair follicle (HF) lineages
- Molecular identification of an asymmetric HF-bulb structure
- Spatial map of fibroblast subtypes in the skin
- Online tool. http://kasperlab.org/tools

## INTRODUCTION

Skin is the largest and one of the most complex organs of the body. A rich variety of epithelial and mesenchymal cell types orchestrate the cyclical hair growth and keep up the barrier function of the skin (Arwert, Hoste, & Watt, 2012; Hsu, Li, & Fuchs, 2014a). For its constant renewal, the skin depends on a range of stem and progenitor cells within its three layers, the epidermis, dermis, and hypodermis, which are either activated steadily or in bouts. Because of this dynamic nature with continuous and cyclical cellular differentiation from stem and progenitor cells towards numerous mature lineages, the mouse skin has become one of the most important model systems to study the genetic and cellular factors that control stem cell plasticity, lineage specification and tissue renewal at large.

However, the full complexity of the skin has not been realized due to insufficient cellular, molecular and spatial characterization of full-thickness skin. To date, the most in-depth single-cell analyses of adult skin, using single-cell RNA-sequencing (scRNA-seq) on selected cell types, collectively revealed for example 25 populations in telogen epidermis, 4 dermal papilla populations and a large variety of fibroblast and anagen HF progenitor populations (Cheng et al., 2018; Ghahramani et al., 2018; Joost et al., 2016; Philippeos et al., 2018; Tabib et al., 2018; Yang et al., 2017). Although most of these studies were restricted by *a priori* defined markers and biased sampling strategies, they uncovered an extraordinary variety of cell types and cell states. However, studies so far have only been scratching the surface of skin cell diversity, leaving major questions unresolved.

How is the skin remodelled to accommodate hair growth? Is this process accomplished through the same cell types that maintain skin at rest, or does it require altered cellular states or even differentiation of new cell types in both epidermis and stroma? The hair follicle (HF) cycles through phases of growth (anagen), regression (catagen) and rest (telogen) (Alonso & Fuchs, 2006; Müller-Röver et al., 2001; Schmidt-Ullrich & Paus, 2005). Hair growth involves rapid expansion of the HF itself to approximately ten times its size, and this process is dependent on remodelling all three skin layers. For example, blood and lymph vessels expand (Mecklenburg et al., 2000), nerves remodel (Botchkarev et al., 1999), immune cell types and numbers change (Castellana, Paus, & Perez-Moreno, 2014), adipocytes mature (Festa et al., 2011), and fibroblast provide new extracellular matrix (Driskell et al., 2013; Rinkevich et al., 2015). However, a systematic investigation of how cell types change in identity and numbers during hair growth has not been performed. The most abundant and best-studied cell type are the keratinocytes (i.e. the epithelial skin cells), which generate the skin barrier by differentiation within the interfollicular epidermis (IFE), and organize into the outer layer, called outer root sheath (ORS), and the seven inner layers of the anagen HF that are generated by the matrix progenitor cells (Langbein & Schweizer, 2005; Legué & Nicolas, 2005; Sequeira & Nicolas, 2012). Until now, the molecular changes that matrix progenitors undergo during inner-layer differentiation are not fully resolved and the composition of the ORS (cell types and molecular profiles) is elusive.

Here, we used scRNA-seq to study the entire repertoire of cells present in full-thickness skin during growth and rest in order to systematically identify their gene expression identity and spatial location. Our unbiased analysis led to several striking discoveries. The ORS is composed of two transcriptionally distinct cell types, each consisting of several subtypes found in specific locations along the HF axis. The HF matrix lineages generating the inner concentric layers branch from transcriptionally uncommitted progenitors, and differentiation of these inner lineages passes through intermediate states that are distinct from their progenitor and end states. The fibroblast subtypes correlate to a spatial axis, and the abundance of two subtypes is hair cycle dependent. Finally, we characterized the companion layer (classically one of the seven inner layers) as transcriptionally belonging to the outer HF layers, and molecularly characterize an asymmetric structure within the HF matrix.

## RESULTS

### Single-cell transcriptome atlas of full-thickness skin during growth and rest

In order to determine the full repertoire of cell types and gene expression programs present in the skin, we isolated cells from the dorsal mouse skin in late anagen (5 weeks old) and telogen (9 weeks old) and generated single-cell transcriptome libraries on the droplet-based 10X Chromium system (**Figure 1A**). We repeated the single-cell sequencing procedure from several independent mice (3 anagen, 2 telogen) and staged each mouse from embedded dorsal skin sections as anagen IV – VI (5-week-old mice) and deep telogen (9-week-old mice) (**Figures S1A-S1B**) (Müller-Röver et al., 2001). To robustly assign cell types, we clustered the single-cell transcriptomes from anagen (3,149 cells) and telogen (2,618 cells) using two parallel approaches: clustering with affinity propagation as previously used in skin (Joost et al., 2016), and graph-based clustering implemented in the standard *Seurat* pipeline (Butler et al., 2018), which were in good agreement (**Figures S1C, S1E-S1F** and **Methods**). Overall, we were able to assign all sequenced skin cells to seven main classes of cells (**Figures 1B, S1H-S1J** and **Tables S1-S2**). While most sequenced cells were keratinocytes derived from either the permanent part of the epidermis (IFE, sebaceous gland and non-cycling portion of the HF, n=2,811) or the cycling part of the anagen HF (n=1,974), we also identified substantial numbers of skin fibroblasts and fibroblast-derived cells (n=385), immune cells (n=210), vascular cells (n=165), melanocytes and Schwann cells (here condensed as neural crest-derived cells, n=144), as well as small numbers of skeletal muscle and red blood cells (n=78, grouped as miscellaneous). Altogether, the main populations could be further separated into 55 transcriptionally distinct subpopulations (listed in **Figure 1B**), reflecting the large complexity of cell types and subtypes present in full-thickness skin.

**Figure 1.**
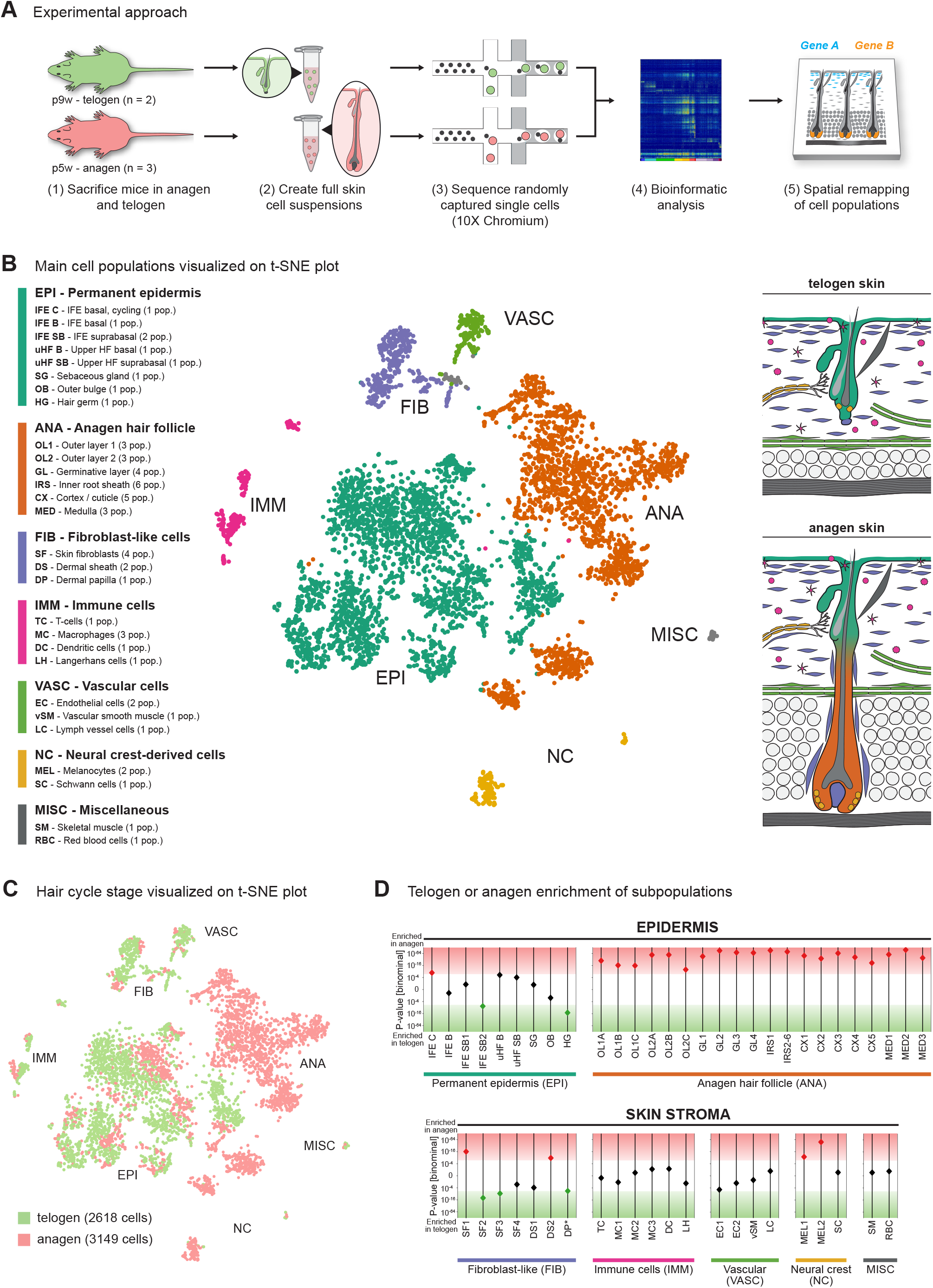
Comprehensive single-cell transcriptome atlas of skin during hair growth and rest. (**A**) Experimental overview. In brief, whole skin cell suspensions were prepared from mice in telogen (p9w, n=2) and anagen (p5w, n=3). Single-cell RNA-seq libraries were generated on the 10X Chromium system, sequenced on a HiSeq2500 and computationally analysed in depth. Selected cell population were then “remapped” to their original spatial position using smRNA-FISH. (**B**) Center: t-SNE visualization of high-quality cells (n=5,767) coloured according to main cell populations. Left: list of distinct cell populations contained within each main population. Right: illustrations representing the microanatomy and the main cellular compartments of the murine skin in telogen (upper) and anagen (lower), with cell colours matching those in the t-SNE. (**C**) t-SNE visualization of high-quality cells coloured according to hair cycle stage. (**D**) Graphical representation of statistical testing for cellular enrichment in telogen or anagen for each cell population independently, grouped according to the main types of cells in the skin. Here, IRS subclusters 2 to 6 are considered as one IRS population (IRS2-6). Asterisk: DP cells from anagen skin were not recovered. P-values were computed using the binomial test and based on absolute cell numbers per skin area.

Next, we determined the contribution of cells sampled from anagen or telogen skin to the different cell types (**Figure 1C**), and investigated whether the cell types and subtypes were maintained at similar cell numbers or statistically enriched in anagen or telogen skin (i.e. occur in higher absolute cell numbers per area) (**Figure S1D** and **Methods**). Reassuringly, most cell populations of the permanent epidermis were present at similar proportions in both anagen and telogen, whereas, as expected, all twenty cell populations derived from the anagen HF and two melanocyte populations were significantly enriched in anagen and the secondary hair germ cells were only identified in telogen samples (P<0.001, Bonferroni-corrected, binomial test) (**Figures 1D** and **S1G**). Interestingly, we identified seven fibroblast-like cell populations (consisting of fibroblasts, dermal sheath and dermal papilla cells), four of which were significantly expanded in either anagen or telogen, demonstrating that the composition of fibroblasts undergoes large-scale remodelling between telogen and anagen (**Figure 1D**). In the following sections, we performed in-depth analyses of selected cell populations to highlight the rich nature of the generated full-thickness skin transcriptome atlas.

### The outer root sheath is composed of two transcriptionally distinct cell types

Several keratinocyte populations were found exclusively in anagen (**Figures 1C-1D**). According to current knowledge we expected to find two major clusters of the cycling part of the anagen HF: one comprising the outer root sheath (ORS; maintaining contact with the basement membrane) and one including the matrix progenitors and the concentric inner layers. However, three main clusters of cells were identified (**Figure 2A**), which were robustly determined using both *Seurat* and AP clustering (**Figures S2A-S2B**). The largest cluster showed a branching topology and was marked by *Msx2* expression (**Figure 2B**), which would be expected for the matrix and inner layers of the anagen HF. The two smaller clusters were characterized by high levels of *Barx2* and *Il11ra1*, respectively (**Figure 2B**). These two clusters have no known counterparts in the literature. We therefore used single-molecule RNA fluorescence *in situ* hybridisation (smRNA-FISH) to spatially map these two cell clusters within the anagen HF, which revealed that they corresponded to spatially distinct parts of the ORS, an ORS-adjacent suprabasal layer, and an internal asymmetric structure (**Figure 2C**). As *Barx2* and *Il11ra1* expression predominantly mapped to the outermost layers of the anagen HF (and had elevated to high *Krt5* and *Krt14* expression) we termed these two major clusters outer layer 1 (OL1; marked by high *Barx2* expression) and outer layer 2 (OL2; marked by high *Il11ra1* expression) (**Figure 2C**). The spatial mapping of the third major cluster of *Msx2*+ cells indeed corresponded to the matrix and inner layers of the anagen follicle (**Figure 2D**), and was further analysed in the next section.

**Figure 2.**
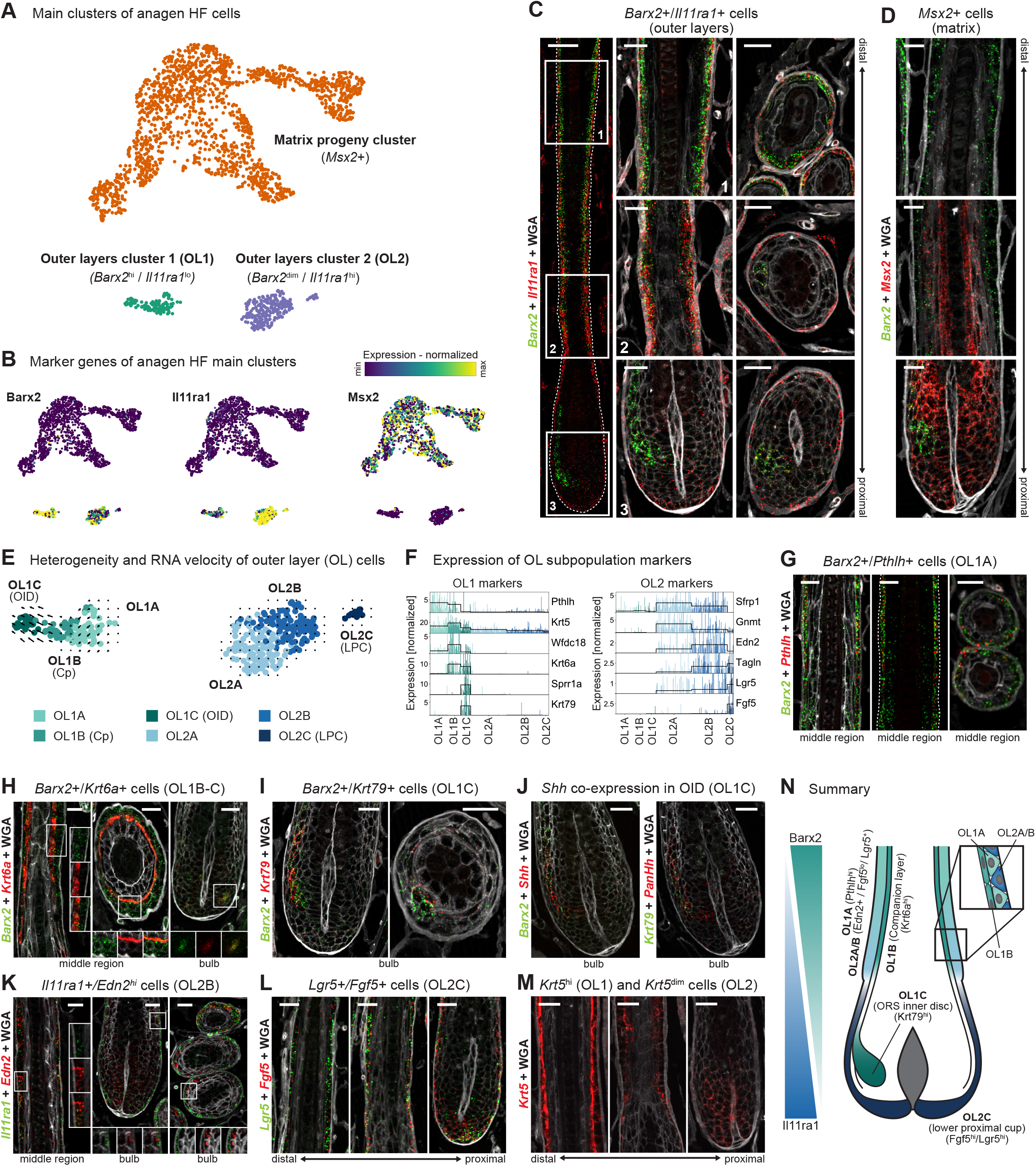
Spatial location and transcriptional signatures of distinct outer root sheath populations. (**A**) Anagen HF cells visualized using UMAP and coloured according to the three main clusters of the anagen HF. (**B**) Expressions of *Barx2*, *Il11ra1* and *Msx2* projected onto the UMAP introduced in (A). (**C**) Spatial location of *Barx2* and *Il11ra1* expressing cells in anagen HF determined by smRNA-FISH. (**D**) Spatial location of *Msx2* expressing cells in anagen HF determined by smRNA-FISH. (**E**) RNA velocity vectors for OL1 and OL2 cells projected onto part of UMAP introduced in (A). (**F**) Expression of marker genes associated with OL1 (left) and OL2 (right) subpopulations. Each bar represents a single-cell transcriptome. (**G-M**) Spatial location of OL subpopulations in anagen HF determined by smRNA-FISH for the following markers: *Barx2* and *Pthlh* expressing OL1A cells (G), *Barx2* and *Krt6a* expressing OL1B-C cells (H), *Barx2* and *Krt79* expressing OL1C cells (I), *Barx2*, *Shh*, *Krt79* and *PanHh* expression in OL1C cells (J), *Il11ra1* and *Edn2* expressing OL2B cells (K), *Lgr5* and *Fgf5* expressing OL2C cells (L), *Krt5^hi^* expressing OL1 cells and *Krt5^dim^* expressing OL2 (M). (**N**) Illustration summarizing the spatial location of molecularly defined outer root sheath cell populations. All smRNA-FISH stainings were performed for n **≥** 3 mice (C-D, G-M). Scale bars: 20 µm (C-D, G-M), 50 µm (C, overview).

Intrigued by our discovery of two transcriptionally distinct types of OL cells, we investigated them in further detail. Importantly, as the OL1 and OL2 cells formed two transcriptionally distinct populations (**Figure 2A**), it suggested that these two cell types are autonomous in the mature anagen follicle (i.e. not connected by intermediate states). To test this hypothesis, we inferred the dynamics of their cellular identities based on the expression patterns of unspliced and spliced mRNAs (RNA velocity) (La Manno et al., 2018). This analysis further corroborated that OL1 and OL2 were distinct cell populations, as RNA velocity did not identify any signs of transition between the two cell populations (**Figures 2E** and **S3A**). Interestingly however, RNA velocity identified a continuous trajectory for the cells within the OL1 cluster, whereas the OL2 cells had no apparent dynamics and represented a more static state (**Figure 2E**).

### The companion layer and an asymmetric inner structure exhibit an OL-cell identity

To drill down on the heterogeneity within the two major OL-cell types, we subclustered the OL1 and OL2 cells, which revealed three subpopulations each (**Figure 2E**). The OL1 subpopulations were characterized by high levels of *Pthlh* (OL1A), *Krt6a* and *Wfdc18* (OL1B), and *Krt79* and *Sprr1a* (OL1C), respectively (**Figures 2F, S2D** and **Table S1**). Based on smRNA-FISH stainings using *Pthlh*, *Barx2*, *Krt6*, and *Krt79*, OL1A cells are located in the basal layer of the ORS (**Figure 2G**), OL1B cells mapped to the companion layer (Cp) (**Figure 2H**), and OL1C cells were located within internal structures (matrix/inner layers) forming an asymmetric structure, which we termed **O**RS **i**nner **d**isc (OID) (**Figure 2I**). Such asymmetric expression pattern was previously described for *Shh* expression and termed the lateral disc (Panteleyev, Jahoda, & Christiano, 2001). Interestingly, the OL1C cells express *Shh* but likely constitute only a portion of the lateral disc, as *Shh* smRNA-FISH staining covered a larger area than the OL1C cells (**Figure 2J**). While the cellular origin of the OID (OL1C) remains elusive, RNA velocity analysis suggests clonal dynamics from OL1A to OL1B to OL1C (**Figures 2E** and **S3A**).

The OL2 subpopulations were characterized by expression of *Edn2* (OL2A and OL2B) and *Fgf5* (OL2C) (**Figures 2F** and **S2D**). OL2A and OL2B were robustly subdivided by different levels of *Col16a1* and *Gnmt* (**Table S2**), however it was not possible to assign their individual locations by RNA staining. *Edn2* smRNA-FISH staining mapped OL2A/B primarily to the basal ORS distal to the bulb (**Figure 2K**), and OL2C (*Fgf5*-expression) is mostly restricted to the lower proximal cup (LPC) (**Figure 2L**). Interestingly, smRNA-FISH staining revealed that the upper and middle part of the ORS showed a particular chessboard pattern of *Krt5*^hi^ cells (likely reduced basement membrane contact of OL1 cells) interspersed with *Krt5*^dim^ cells (likely increased basement membrane contact of OL2 cells), suggesting that OL1 and OL2 cells intermix in parts of the ORS (**Figure 2M**). Altogether, we discovered six populations of cells with an “ORS-like” transcriptional identity, three of which constitute distinct parts of the ORS, one corresponds to the Cp, one to the LPC and one extending into the internal layers of the HF and constituting parts of the lateral disc (**Figure 2N**).

### Hair follicle matrix lineages branch from transcriptionally uncommitted progenitors

In contrast to the two discrete OL (outer layer) clusters, *Msx2^+^* matrix cells showed the characteristic topology of different lineages branching from a central population of progenitor cells (**Figures 3A** and **S2B**). The anagen HF is known for its distinctive microanatomy composed of seven concentric inner layers: the companion layer (Cp), three layers of the inner root sheath (IRS: Henle (He), Huxley (Hu), IRS cuticle (Icu)), and the cuticle (CU), cortex (CX) and medulla (MED) layers that form the new hair shaft (Fuchs, 2007). These layers are maintained by the germinative layer (GL) cells, which are located adjacent to the dermal papilla. Importantly, marker gene expression defined in this study (**Figures 3B, S2D** and **Table S1**) and classification based on keratin expression patterns in human skin (Langbein et al., 2010) (**Figure S2F**) both supported a central core of GL cells and three branches that represent differentiation into the IRS, cortex / cuticle (CX), and MED layers. The GL cells were demarcated by high expression of *Mt2* and *Dcn*, and based on their gradually changing expression levels of genes such as *Id3* and *Lef1* (DasGupta & Fuchs, 1999; Genander et al., 2014) GL cells separated into four populations (GL1-4) (**Figures 3A-3B** and **Table S1**).

**Figure 3.**
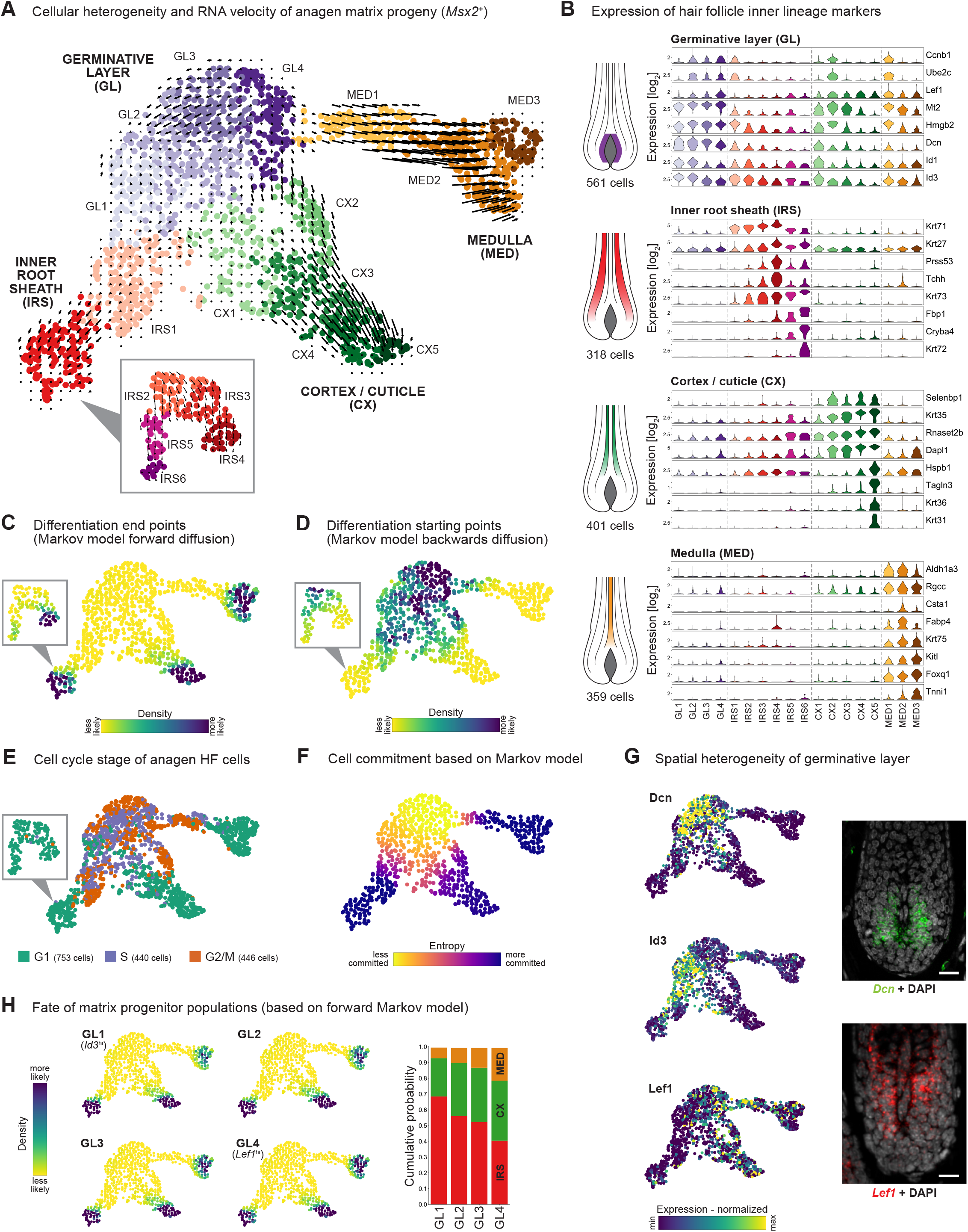
Reconstruction of inner-layer differentiation in the anagen hair follicle. (**A**) RNA velocity vectors for *Msx2+* cells, coloured according to subclusters (GL, IRS, CX and MED) and projected onto part of the UMAP introduced in Figure 2A. Inset: subclustering of differentiated IRS cells reveals two trajectories. (**B**) Violin plots showing expressions of markers for all the subpopulations. (**C-D**) Differentiation end points (C) and starting points (D) inferred using a Markov model based on RNA velocity data introduced in (3A). Inset data is based on a separate Markov model including only IRS2-6 cells. (**E**) Inferred cell cycle stage for cells displayed in the UMAP introduced in (3A). (**F**) Quantification of cell commitment for each independent cell, based on Markov processes proceeding from each cell, displayed on the UMAP introduced in (3A). (**G**) Spatial location of GL cells. Left panels: expression of *Dcn*, *Id3* and *Lef1* projected onto part of the UMAP introduced in (2A). Right panels: Spatial expression of *Dcn* and *Lef1* in germinative layer of anagen HF mapped using smRNA-FISH (n = 3 mice). (**H**) Left: Quantification of fate bias for independent cells of the four GL subtypes based on Markov processes proceeding from each cell, displayed onto part of the UMAP introduced in (2A). Right: Barplot that summarizes the cell fates of cells belonging to each GL subtype. Scale bars: 20 µm (G).

For the three branches of inner HF lineages, we could divide the cells of each branch into several subpopulations, which represent early, intermediary or mature stages of the respective differentiation process. For the IRS lineage, an “early” cell cluster (IRS1) co-expressed both IRS and GL markers, whereas the more differentiated cells (IRS2-6) could be further separated along two branches (inset in **Figure 3A**) by a combination of known and novel genes (**Figures 3B, S2E** and **Table S1**). For example, the IRS3/IRS4 branch expressed *Krt71* and *Tchh* at high levels (Fuchs, 2007) as well as the protease *Prss53* (**Figure 3B**), which has been previously shown to influence hair shape in a genome wide association study (Adhikari et al., 2016). In contrast, the IRS5/IRS6 branch showed a more distinct Icu identity marked by *Krt72* and *Krt73* expression (Langbein et al., 2010), the metabolic enzyme *Fbp1* and the crystallin *Cryba4* (**Figures 3B**). Although we successfully distinguished the Icu branch, it is unclear to what proportion the other branch (IRS3/4) contains Henle layer and Huxley layer cells. Similar to the IRS, cells associated with CX differentiation could be divided into distinct stages marked by increasing expression of *Krt35* (Langbein et al., 2010), the ribonuclease *Rnaset2b* and the differentiation associated protein *Dapl1* (**Figure 3B**). While stage CX4 is compatible with both a cortex and cuticle identity (**Figure S2F**), stage CX5 is expressing cortex specific markers such *Krt31*, *Krt36* and actin-linking protein *Tagln3* (**Figures 3B** and **S2D**). Finally, the MED cells were divided into three stages, corresponding to an early differentiation stage (MED1) co-expressing low levels of MED markers with GL and cell cycle-associated genes, and two mature stages showing high expression levels of the MED marker *Foxq1* (Potter et al., 2006), but diverging in expression of the cysteine protease inhibitor *Csta1*, the fatty acid carrier *Fabp4*, the Kit receptor ligand *Kitl* and the contractile protein *Tnni1* (**Figures 3B** and **S2D**). Interestingly and as previously described (Morioka, 2005), MED cells also expressed *Krt75*, which has previously been defined as a Cp marker (**Figures 3B** and **S2G**) (Sequeira & Nicolas, 2012).

Having used known marker gene expression to validate our unbiased transcriptional reconstruction of the multi-branching differentiation process within the inner HF layers, we next investigated the extent of fate bias among the GL progenitors, and determined when cells commit to the three distinct lineages. Importantly, analysis of RNA velocity confirmed that our matrix subpopulations (GL1-4) represent stages in an active differentiation process towards the inner lineages of the HF, with the field-normalized arrows clearly emerging from GL cells towards the three main branches (**Figures 3A** and **S3B**). Furthermore, an unsupervised Markov model (based on RNA velocity data) predicted that the populations IRS4/6, CX5 and MED2/3 are the endpoints of the differentiation processes, and identified the GL cells as the starting point (**Figures 3C-3D**). The multi-branching topology of uncommitted GL cells towards each lineage was further supported by a shift from proliferative to post-mitotic cellular identities of cells within each separate branch (**Figure 3E** and **S2C**). RNA velocity analyses also indicated that lineage commitments first emerge in cells located at the root of each branch (IRS1, CX1/2, MED1), while cells in the GL compartment (GL1-4) are still uncommitted (**Figure 3F**). Following up on GL-cell commitment, GL1-4 cells showed differential expression of certain genes along the dermal papilla axis (e.g. *Id3* and *Lef1*) pointing towards one of the three branch endpoints (**Figure 3G**), which may indicate a potential fate bias. The Markov model indeed predicted a small but not significant bias for GL1 toward IRS and GL4 towards CX and MED; however, each GL subpopulation (GL1-4) can clearly give rise to all branch endpoints (**Figures 3H** and **S3C-S3D**). Importantly, a recent study used single-cell RNA-sequencing data of sorted matrix cells to describe that spatially defined micro-niches (i.e. dermal papilla cells juxtaposed to individual GL cells) regulate cell fate (i.e. determine the GL progenitor differentiation path) (Yang et al., 2017). Interestingly, comparison of these marker-based FACS sorted matrix cells to our anagen HF transcriptomes revealed that only a subset represents GL cells whereas many cells transcriptionally matched intermediate or terminally differentiated cell identities (**Figure S3E**). While it is very likely that differentiation of a GL cell into a specific lineage of IRS, CX or MED is dependent on its exact position (influenced by micro-niche signalling), our data supports that cells from all GL populations are transcriptionally uncommitted and do not express signatures of any branch.

### Differentiation of inner layers transits through intermediate molecular programs

Having established that the *Msx2*-expressing cells captured the differentiation processes from uncommitted GL cells to differentiated cells of the inner layers of the HF, we next sought to identify the expression programs induced as the cells committed and differentiated. By modelling the data as a diffusion process (Haghverdi et al., 2016), we arranged the cells on the IRS, CX and MED branches in a pseudotemporal order (**Figures 4A** and **S4A**). This facilitated the identification of around 1000 genes each that significantly change in expression along the IRS, CX and MED branch, respectively, which were further grouped based on their co-expression patterns into modules. All branches had in common that the first four modules were characterized by GL-related genes that become downregulated during the differentiation processes (**Figures 4B** and **S4C**). However, inner layers differentiation was not just consisting of the gradual downregulation of GL genes and the simultaneous upregulation of terminal inner lineage signatures. Instead, cells of all lineages passed through distinct intermediate states marked by modules of genes (V-VI) found neither in the uncommitted progenitor population, nor in the terminal differentiated state (**Figure 4B**). To illustrate genes that were expressed transiently, we visualized *Ctsc*, *Selenbp1* and *Aldh1a3* expression along the respective branch (**Figure 4C**). These intermediate populations were also associated with the shift from proliferative to post-mitotic cell identity (**Figures 3E** and **4B**), the acquisition of a deterministic cell fate (**Figure 3F**), and their lineage-associated differentiation markers (**Figures 4D** and **S4D-S4E**).

**Figure 4.**
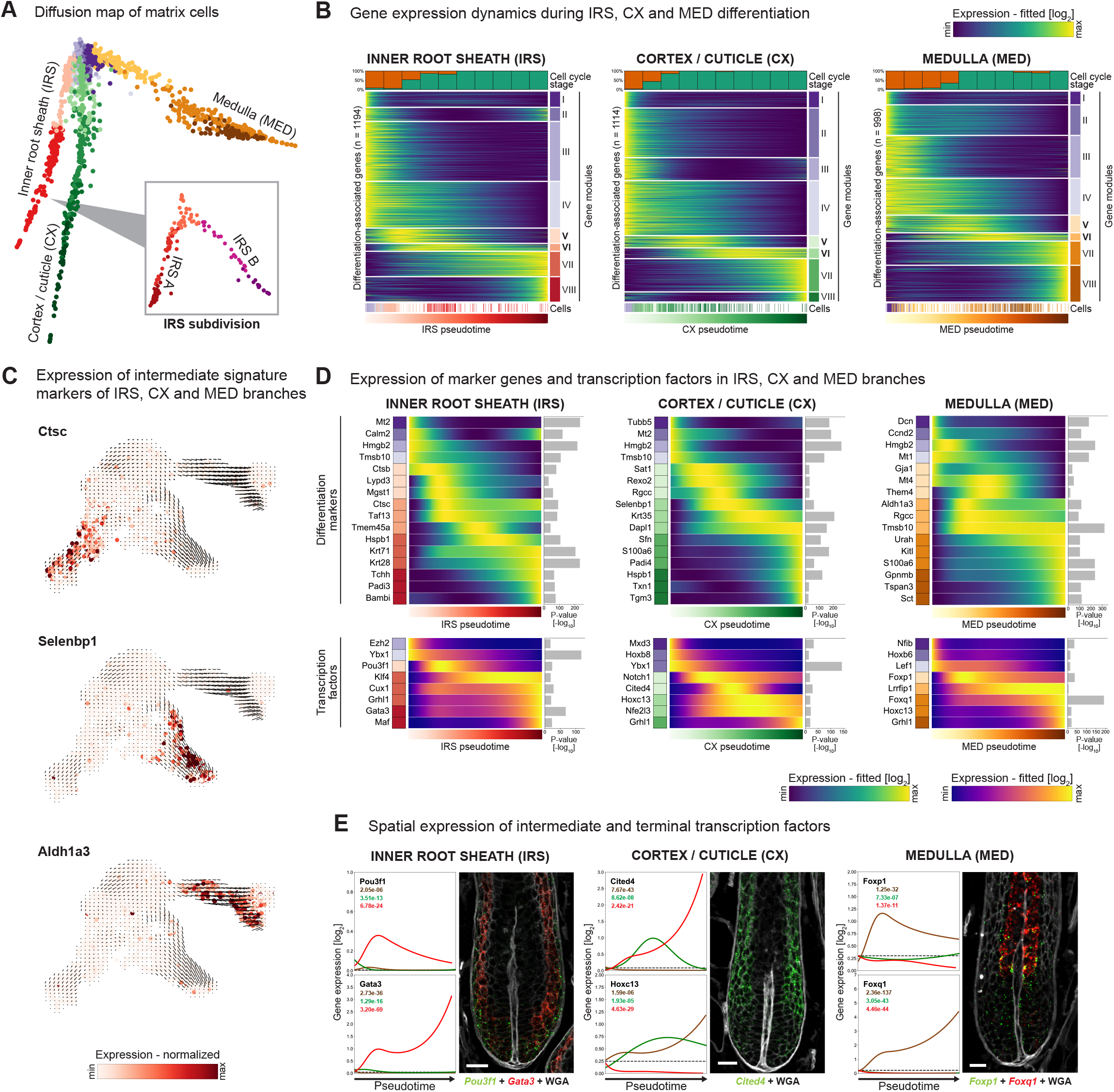
Intermediate and terminal gene expression signatures during inner-layer differentiation. (**A**) Anagen HF matrix progeny projected onto diffusion map. Inset: IRS subpopulations projected onto diffusion map of IRS cells. (**B**) Heatmaps showing genes (fitted expression) with significant association to lineage-differentiation of IRS (1194 genes), CX (1114 genes) or MED (998 genes), respectively. Genes are grouped into modules (right panel) and individual cells are positioned along diffusion pseudotime (lower panel). The upper panel shows the cell cycle stage (green: G1; red: S/M/G2) of cells binned along pseudotime. (**C**) Expression of the transiently-upregulated markers *Ctcs*, *Selenbp1* and *Aldh1a3* overlaid onto RNA velocity vectors introduced in Figure 3A. (**D**) Heatmaps exemplifying genes with significant lineage-associated expression patterns (fitted), along the pseudotime of each lineage separately. Upper: Marker genes that become downregulated, transiently upregulated or permanently upregulated during inner layer differentiation. Bottom: Heatmap with transcription factor expressions (fitted) along the differentiation pseudotime of each lineage. (**E**) Left panels: comparative expression (fitted) of transcription factors in IRS (red), CX (green) and MED (brown) branches. Dotted lines show the null model (baseline expression in root cell cluster GL4). Numbers denote the P-values for branch-specific expression dynamics (both increased and decreased expression) compared to a reduced model which assumes equal expression over all branches. Right panels: Spatial positioning of cells expressing selected transcription factors by smRNA-FISH (n = 3 mice). Scale bars: 20 µm (E).

To investigate the regulation of lineage-specific gene expression programs, we identified transcription factors with significant lineage-associated expression (**Figures 4D** and **S4F**). GL cells expressed *Ybx1* and the homeobox protein *Hoxb6* (**Figure 4D** and **Table S1**). Intermediate cell identities expressed distinct transcription factors, exemplified by expression of *Pou3f1* in IRS cells (and in the LPC) and *Foxp1* in MED cells (**Figure 4E**). Intermediate CX-lineage cells were not associated with the expression of a unique transcription factor but rather shared upregulation of *Cited4* with IRS cells (**Figure 4E**). Spatial mapping of intermediate transcription factors indicates that their expression peaks in the matrix outside the GL (**Figure 4E**). Transcription factors identified in more terminally differentiated cells included well-known markers such as *Cux1*, *Gata3* and *Maf* for IRS specification (Ellis, 2001; Kaufman et al., 2003), and *Foxq1* for MED differentiation (**Figures 4D-4E** and **S4F**) (Potter et al., 2006). Again, we did not find a transcription factor that was specific for terminally differentiating CX cells, as *Hoxc13* expression was shared with the MED branch (Godwin & Capecchi, 1998) (**Figures 4D-4E**). Interestingly, we identified several additional transcription factors that were induced in two or more branches, most importantly *Grhl1*, which was associated with terminal differentiation in all three branches (**Figure S4F**), in addition to its known dynamical expression during IFE differentiation (Joost et al., 2016). Noteworthy, analysis of the IRS subbranches, IRS A and IRS B (**Figures 4A, S4B** and **S4G-S4H**), revealed expression of the transcription factors *Maf* and *Mafb* within the IRS A branch (**Figure S4I**). Altogether, our analysis identified lineage-specific expression of several transcription factors that have not been implicated in inner layer differentiation before, while taking its credibility from the confirmation of known transcription factors.

Overall, the analyses of the anagen HF inner-layer differentiation at single-cell resolution revealed a multistep process. Cells exit the cell cycle in conjunction with the induction of an intermediate molecular program that ensures cell differentiation along a specific lineage, with the ultimate specification of a terminally differentiated cell identity.

### Transcriptionally distinct fibroblasts populate different layers of the skin

Having analysed gene expression and cellular heterogeneity of the epithelial compartment, we next focused on fibroblast and fibroblast-like cells as epidermal keratinocytes closely interact with these populations. Although a variety of fibroblast subtypes have been defined in embryonic and adult skin, the full extent of fibroblast heterogeneity in the skin is not resolved (Driskell et al., 2013; Korosec et al., 2019; Philippeos et al., 2018; Rinkevich et al., 2015, Tabib et al., 2018). Overall, our analysis separated the fibroblasts and fibroblast-like cells into seven cell populations (**Figures 5A, S5A** and **Table S1**). These corresponded to four skin fibroblast subtypes (SF1-4), dermal papilla cells (DP), mature dermal sheath cells (DS2), and one with an intermediary signature between SF and DS (DS1) (**Figures 5A-5B** and **S5A-S5B**). Interestingly, the skin fibroblast and fibroblast-like populations did not form discrete clouds but rather formed a connected structure that suggested gradually changing cellular identities across these populations (**Figure 5A**). Reassuringly, this topology was reproducible using both UMAP and t-SNE (**Figures 5A** and **S5B**). Moreover, the fibroblast populations showed high plasticity between telogen and anagen skin (**Figure 1D**), with SF1 being highly enriched and SF2 being completely depleted in anagen skin (**Figures 5C** and **S5C**). Also, DS2 was significantly enriched in anagen, whereas DS1 was enriched in telogen skin (**Figures 1D** and **S5C**).

**Figure 5.**
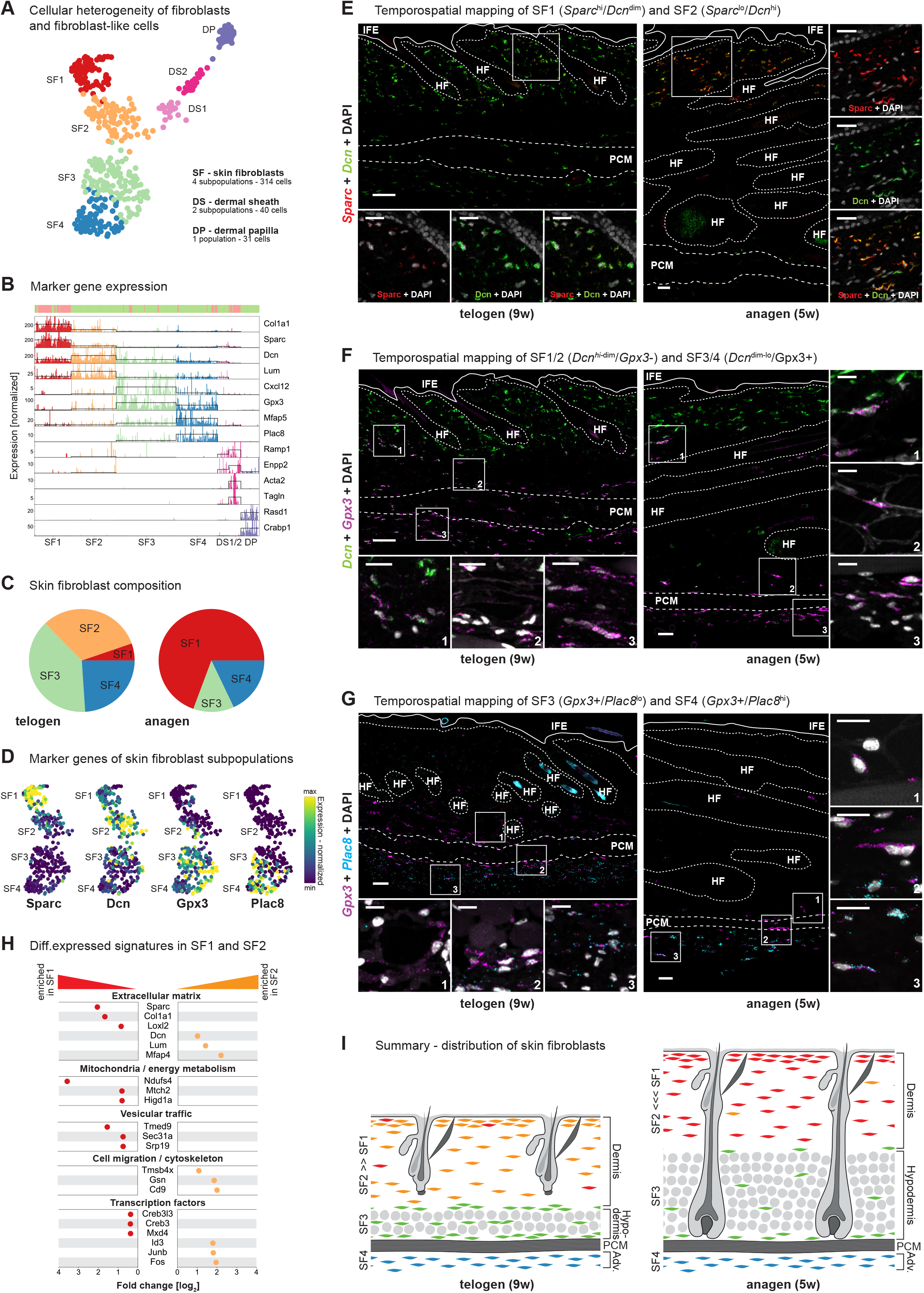
Transcriptional heterogeneity and spatial location of fibroblasts and fibroblast-like cells. (**A**) Subclusters of fibroblasts and fibroblast-like cells projected onto UMAP representation of fibroblasts and fibroblast-like cells. (**B**) Expression of marker genes associated with each fibroblast and fibroblast-like population. Each bar represents a single-cell transcriptome. Hair cycle stages of cells are shown in the top panel (green: telogen; red: anagen). (**C**) Distribution of fibroblast subtypes in anagen and telogen skin. (**D**) Expression of fibroblast subtype markers *Sparc*, *Dcn*, *Gpx3* and *Plac8* projected onto part of UMAP introduced in (A). (**E-G**) Temporo-spatial mapping of skin fibroblast subtypes in anagen and telogen skin using smRNA-FISH. Images show both overviews of skin, and detailed zoom-ins complemented with nuclear stain. HF: Hair follicle. IFE: Interfollicular epidermis. PCM: *Panniculus carnosus* muscle. (E) smRNA-FISH against Sparc and Dcn that distinguish SF1 and SF2 subtypes of fibroblasts. (F) smRNA-FISH against Dcn and Gpx3 that distinguish SF1/2 and SF3/4 subtypes of fibroblasts. (G) smRNA-FISH against Gpx3 and Plac8 that distinguish SF3 and SF4 subtypes of fibroblasts. (**H**) Differential expression of selected genes between SF1 (enriched in anagen) and SF2 (enriched in telogen), shown with fold-changes. All genes had significant differential expression with adjusted P-value < 0.05. (**I**) Illustration summarizing the spatial distribution of skin fibroblasts during anagen and telogen. All smRNA-FISH stainings were performed for n **≥** 3 anagen mice and n **≥** 2 telogen mice (E-G). Scale bars: 50 µm (overviews), 20 µm (insets) (E-G).

To investigate the spatial distribution of the fibroblasts (SF1-4), we identified four marker genes (*Dcn, Gpx3, Sparc* and *Plac8*) that together separated these fibroblast subtypes (**Figure 5D** and **S5D**). smRNA-FISH for *Dcn*-expressing SF1 and SF2 cells revealed that they are located in both the papillary and reticular dermis (**Figure 5E**), while the *Gpx3*-expressing fibroblasts (SF3 and SF4) are distributed in the hypodermis, around the *Panniculus carnosus* muscle and in the adventitia (**Figure 5F**). smRNA-FISH of *Plac8* revealed the location of SF3 (*Gpx3*+/*Plac8*-) cells in the hypodermis and around the *Panniculus carnosus* muscle, while SF4 cells (*Gpx3*+/*Plac8*+/*Mfap5*+) were located in the adventitia (**Figures 5G** and **S5E**). Interestingly, we observed a striking difference in *Sparc* expression that distinguished anagen SF1 (*Sparc*^hi^/*Dcn*^dim^) from telogen SF2 (*Sparc*^dim^/*Dcn*^hi^). SPARC is a matricellular protein (i.e. a dynamically expressed glycoprotein modulating extracellular matrix interactions) and commonly overexpressed in fibrotic diseases (Wang et al., 2010). According to our data, *Sparc* is only lowly expressed during the telogen phase, the predominant hair cycle stage in mammalian skin. Anagen SF1 compared to telogen SF2 fibroblasts also displayed increased expression of genes linked to collagen production and associated processes such as vesicular trafficking and energy metabolism. SF1 cells are furthermore distinguished by the induction of transcription factors such as *Creb3*, *Creb3l3* and *Mxd4*. SF2 cells express an ECM signature dominated by proteoglycans such as *Dcn*, *Lum* and *Mfap4*, as well as genes linked to increased migration and cytoskeletal remodelling (**Figure 5H**).

As predicted by our single-cell transcriptome data (**Figure 5D**), we indeed observed cells expressing mixed signatures at the interface of each fibroblast compartment supporting a gradual change of cellular identities across fibroblast subtypes (**Figures 5E-5G**). Overall, we unveiled previously unknown transcriptional heterogeneity among skin fibroblasts, revealed their respective locations in the different skin layers and defined new markers for these fibroblast subtypes. In addition, we observed a striking remodelling of dermal fibroblast in association with the hair cycle stages, summarized in **Figure 5I**.

### Minor transcriptional changes in the permanent epidermis, immune cells, vascular cells and neural-crest derived cells between hair cycle stages

Detailed analysis of the cells belonging to the permanent part of the epidermis demonstrated that all subpopulations resolved in this study corresponded to those defined in our previous study (Joost et al., 2016). This includes subpopulations of the interfollicular epidermis (IFE), upper HF, sebaceous gland and outer bulge (**Figures 6A**, **S6A-S6B, S6I** and **Table S1**). Because of differences in the cell capturing approach (10X Chromium versus Fluidigm C1 system) or the adapted protocol for full-thickness skin cell sampling, we did not capture inner bulge and cornified layer keratinocyte in this study, which were analysed earlier (Joost et al., 2016). However, we now resolved bulge-like *Cd34*-negative and *Id3*-positive cells likely corresponding to the secondary hair germ of the telogen HF (Genander et al., 2014) (**Figure 6A**). All permanent epidermis cell populations were sampled from both anagen and telogen skin without enrichment towards any hair cycle stage (**Figures 1D** and **S6C**). Nevertheless, we identified limited but robust transcriptional alterations in these cells related to the hair cycle (**Figures S6D-S6E**). In anagen, most of the induced genes (compared to telogen) were involved in metabolic processes, including intermediate metabolism (*Oat, Pkm, Tpi1, Dbi1*), energy metabolism (*Ndufa4l2, Ucp2*), mitochondrial function (*mt-Cytb*), and protein synthesis. This more active cell state in anagen was further supported by observing a higher fraction of proliferating cells in the IFE and the upper HF in anagen skin in line with recent work (Reichenbach et al., 2018) (**Figures S6G-S6H**). In contrast, genes upregulated in telogen included metallothionins (*Mt1* and *Mt2*), and differentiation markers, such as *Sfn* (**Figure S6E**). In the anagen bulge, we additionally found upregulation of genes which were highly induced in the cycling part of the anagen HF (such as *Barx2, Dbi*, and *Ucp2)*, and downregulation of typical telogen bulge markers (for example *Cd34* and *Ftl1)* (**Figures S6E-S6F**). Overall, we observed only subtle gene-expression changes in the permanent epidermis populations.

**Figure 6.**
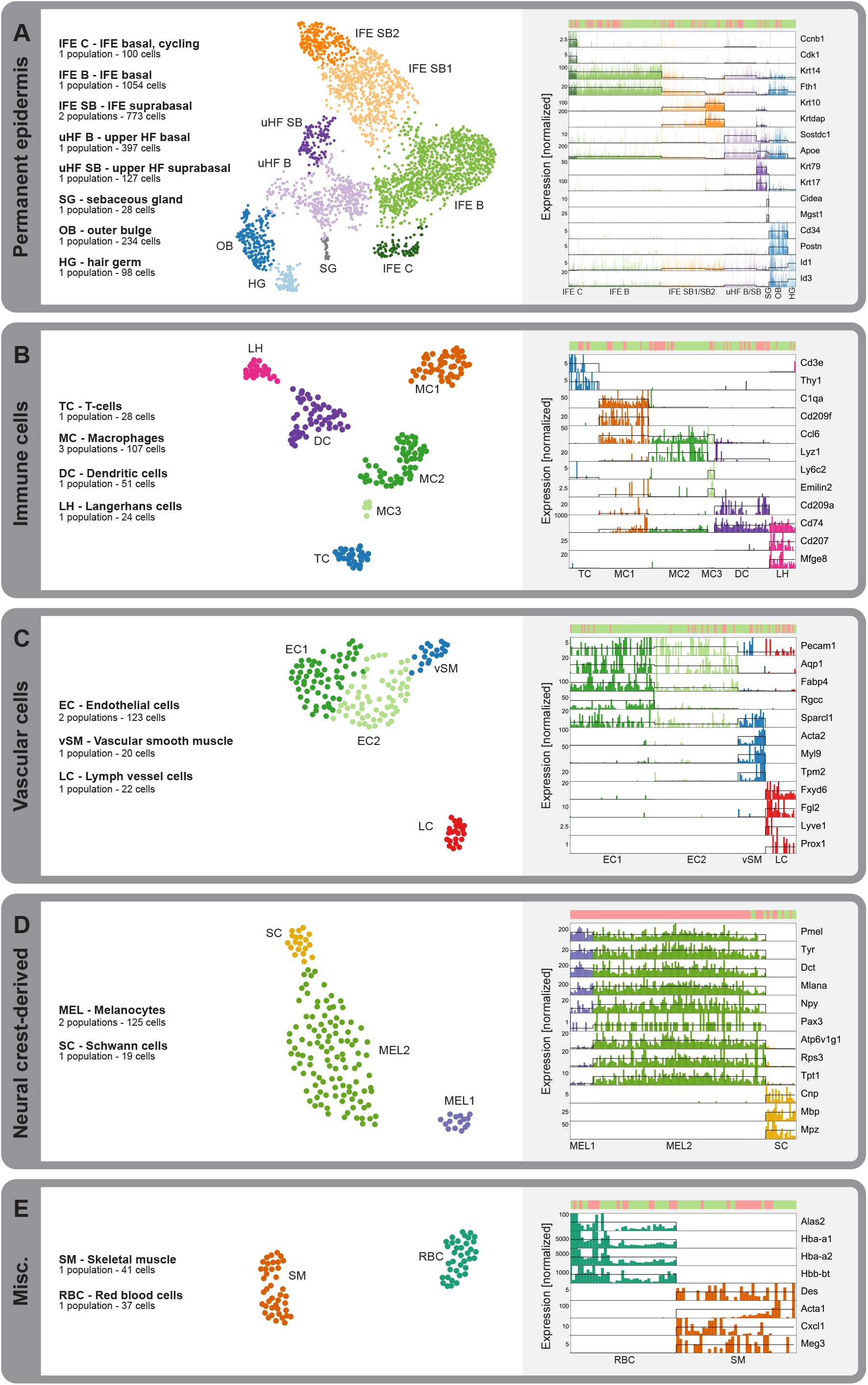
Subclustering of permanent epidermis and remaining stromal compartments. **(A-E)** Subclustering of (A) permanent epidermis, (B) immune, (C) vascular, (D) neural crest-derived and (E) miscellaneous cell populations. Left panels: 2nd level clusters projected onto UMAPs. Right: Expression of marker genes associated with each subpopulation. Each bar represents a single-cell transcriptome. Hair cycle stage of cells are shown in the top panel (green: telogen, red: anagen).

Finally, our single-cell atlas contained diverse other cell types such as immune, vascular, and neural crest-derived cell populations (**Figures 6B-6E, S7A-S7H** and **Tables S1-S2**). Overall, we identified six populations of immune cells in our dataset: T-cells (TC; most likely γδ T-cells) defined by consensus markers such as *Cd3e*, *Thy1* and *Nkg7*, dendritic cells (DC) marked by *Cd209a* and *Cd74*, Langerhans cells (LC) marked by *Cd207* and *Mfeg8*, and three subpopulations of macrophages (MC) characterized by, for example, *Ccl6, C1qa* and *Cd209f* (MC1), *Ccl6* and *Lyz1* (MC2), and *Ccl6, Ly6c2* and *Emilin2* (MC3) (Heng et al., 2008) (**Figures 6B** and **S7M**). None of these six major clusters of immune-cell populations showed clear statistical enrichment in either anagen or telogen skin, suggesting that the repertoire of the major classes of immune cells is mostly unchanged between hair cycle phases (**Figures 1D** and **S7I**). Notably, it has been reported that subtypes of immune cells change during the hair cycle (Castellana et al., 2014), a change that our cell-sampling depth may not be able to pick up. The vascular cells also showed unchanged abundance in anagen and telogen skin (**Figures 1D** and **S7J**). In addition to a lymph vessel cell (LC) population (characterized by *Lyve1*, *Prox1, Fxyd6*, and *Fgl2* expression), we identified vascular smooth muscle cells (vSM) expressing *Sparcl1*, *Acta2, Myl9 and Tpm2*, and two subpopulations of endothelial cells (EC) distinguished by different levels of genes such as *Fabp4*, *Ly6c1* and *Rgcc* (Zhao et al., 2018) (**Figures 6C** and **S7N**). Among neural-crest derived cells, the Schwann cells (SC), which were identified by *Mbp* and *Mpz* expression, showed no hair cycle enrichment, whereas the melanocytes were nearly exclusively found in anagen skin, as expected (Lin & Fisher, 2007) (**Figures 1D** and **S7K**). Interestingly, melanocytes could also be divided into a metabolically more quiescent (MEL1) and a metabolically more active subtype (MEL2) (**Figures 6D** and **S7O**). Finally, we found a population of skeletal muscle cells – likely belonging to the *Panniculus carnosus* – marked by *Des* and *Acta1*, and a population of red blood cells marked by typical genes such as *Hba-a1* (**Figures 6E** and **S7P)**. Neither population showed a clear association to either anagen or telogen skin (**Figures 1D** and **S7L**).

## DISCUSSION

For this study, we generated single-cell transcriptomes from mouse skin during hair growth and rest to systematically investigate the presence and plasticity of cell types and subtypes, together with their respective gene expression programs and spatial positions. We chose the most expanded anagen (IV – VI, 5 weeks) and deep telogen (9 weeks) (Müller-Röver et al., 2001) to define robust end states of differentiation (as opposed to studying the remodelling of the tissue during anagen initiation and expansion), and to study the ongoing differentiation processes of the inner lineages in the mature anagen HF. Overall, we demarcated 55 robust cell populations sampled from nearly all cellular compartments of the skin (**Figure 1**), which led to several important discoveries that we discuss in the following sections.

### Six different cell populations with outer-layer signatures constitute the ORS, LPC, Cp and OID

Our data revealed that the anagen-specific HF cells were highly heterogeneous. They were composed of more than 20 distinct subpopulations that unexpectedly grouped into three, instead of two, main cell types: two distinct ORS-associated types (OL1 and OL2, each containing three subpopulations) (**Figure 2**), and the branching progeny of the HF matrix (**Figure 3**). Using smRNA-FISH, we revealed the spatial organization of the ORS and an uncharacterized asymmetric structure. Rather unexpected, cells of both OL1 and OL2 types were typically intermixed along most of the ORS (OL1A and OL2A/B), with OL2A/B cells having more access to the basement membrane than OL1A cells. Moreover, the OL1 populations (OL1A, OL1B (Cp), OL1C (OID)) were reminiscent to the telogen upper HF, while OL2 populations (OL2A/B and OL2C (LPC)) were distinguished by expression of typical telogen bulge marker genes. The next surprise was that the companion layer (Cp) was found among an OL cluster (OL1B). The Cp separates the ORS and IRS, and is classically described as one of the seven concentric inner layers of the anagen HF (Fuchs, 2007; Rompolas et al., 2013). At anagen onset, the Cp is generated from Krt79-expressing progenitors originated from the secondary hair germ (Veniaminova et al., 2013). During anagen HF expansion, the Cp and ORS also have different cellular origins as the Cp is generated by progenitors located juxtaposed the DP (Mesler et al., 2017; Yang et al., 2017) and the ORS by downward expansion of upper ORS cells (Hsu, Li, & Fuchs, 2014b; Hsu, Pasolli, & Fuchs, 2011). Thus, it was unexpected to find that the Cp was transcriptionally more similar to ORS than inner layer cells. Furthermore unexpected, we identified a column of suprabasal cells extending from the Cp into the IRS/matrix area based on OL markers. These cells expressed for example *Krt5*, *Krt6a* and *Barx2*, and were distinguished by high levels of *Krt79* (OL1C). The morphology of this structure was reminiscent of the asymmetric expression pattern of Sonic Hedgehog (*Shh*) in the HF bulb, which has been previously described as the lateral disc (Gat, et al., 1998; Panteleyev et al., 2001). Our OID structure was indeed partially overlapping with the *Shh-*expressing cells, indicating unique niche signals in this asymmetric area of the hair bulb. The lateral disc cells were originally hypothesized to generate the new secondary hair germ during catagen (Panteleyev et al., 2001), however recent studies demonstrated through lineage tracing that *Shh*-expressing cells instead generate IRS lineages and their traced progeny does not survive catagen (i.e. is not the origin of the secondary hair germ) (Mesler et al., 2017; Panteleyev, 2018; Xin et al., 2018). The transcriptomes of the OID-forming cells did not however cluster together with the IRS populations in our data. Instead the OID-forming population of cells seemed to represent the endpoint of a gradual identity change among OL1 cells (**Figure 2E**). Thus, the functional role of this cell population is not apparent from our data and will be exciting to investigate in future studies.

### Matrix progenitors are transcriptionally uncommitted

Detailed analyses of the *Msx2*+ matrix cells revealed a unified pool of germinative layer cells (GL) that branch into inner root sheath (IRS), cortex / cuticle (CX) and medulla (MED) cells (**Figure 4**). A recent study based on single-cell RNA-sequencing data of sorted matrix cells concluded that spatially defined micro-niches along the proximal-distal axis of the dermal papilla regulate the cell fate of individual GL cells (Yang et al., 2017). While it is very likely that differentiation of a GL cell into a specific inner lineage is dependent on its exact position (and influenced by micro-niche signalling), our data demonstrates that cells from all GL populations are transcriptionally uncommitted and do not express signatures of any branch. We indeed detected some cells that co-expressed a GL signature with branch-specific genes similar to those described by (Yang et al., 2017). However, these cells were located in between the large group of uncommitted progenitors and the respective lineage branches, and thus likely represent the initial differentiation stages of the inner layers. Importantly, our findings do not exclude the presence of micro-niches along the dermal papilla axis (Yang et al., 2017), nor contradict the well-established observation that the spatial position of GL cells influences lineage fate (Legué & Nicolas, 2005; Legué, Sequeira, & Nicolas, 2010; Sequeira & Nicolas, 2012). Our data however proves that these influencing factors are not yet deterministically reflected in the transcriptomes of GL cells. This is significant as a low degree of transcriptional commitment is likely linked to a higher degree of plasticity among GL cells, allowing them to rapidly adapt to changes in the HF environment such as cell loss in the GL compartment. Indeed, a recent study based on spatio-temporal tracking in live mice showed that GL cells flexibly change their position along the proximal-distal axis, and with it their lineage commitment, even in the absence of tissue damage (Xin et al., 2018).

### Inner-lineage differentiations pass through intermediate states

Analysis of RNA velocity clearly showed an active differentiation process starting from the GL and terminating at the end of the IRS, CX and MED branches. Interestingly however, all three branches showed a very similar expression dynamics during differentiation: instead of just up-regulating branch-specific differentiation genes while down-regulating GL signatures, cells on all branches passed through unique intermediate states. These states were characterized by branch-specific gene expression signatures that were distinct from their progenitor and end states (**Figure 4**), and as soon as cells entered any of the intermediate states they stopped proliferating and started to rapidly adapt their transcriptome towards terminal differentiation. Therefore, these intermediate states may mark the *points of no return* in lineage commitment. Also noteworthy, while we found a variety of transcription factors uniquely expressed in either the IRS branch or MED branch, we did not identify transcription factors that were specifically linked to the CX branch (located between IRS and MED). Instead, most transcription factors upregulated in the CX branch were co-expressed in either the MED or the IRS branch. One intriguing hypothesis for this observation could be that CX identity is the default lineage identity of differentiating matrix cells, which needs to be actively directed towards IRS or MED fate by additional transcriptional signatures.

### Systematic comparison of telogen and anagen reveals new insights in the skin stroma

Systematically profiling cells in both telogen and anagen skin enabled us to investigate hair stage-related changes in abundance of cell types and their transcriptional differences. Only modest changes in gene expression were found in the permanent parts of the HF and IFE, and these differences were mainly related to increased metabolism and proliferation in anagen cells, in agreement with recent data describing a transition to higher protein synthesis in the HF bulge compartment during anagen (Blanco et al., 2016). Interestingly, we found that stromal components were dramatically remodelled in skin during hair growth and rest. We demonstrated that two different fibroblast subtypes (SF1 and SF2) occupy the same space in the dermis during telogen and anagen, but SF1 was present exclusively in anagen and SF2 mainly in telogen. The anagen SF1 cells had increased expression of genes linked to collagen production and associated processes such as vesicular trafficking and energy metabolism, whereas SF2 cells expressed an ECM signature dominated by proteoglycans, as well as genes linked to increased migration and cytoskeletal remodelling (**Figure 5H**). It is tempting to speculate that upon the induction of hair growth, SF1 fibroblasts swap identity to become SF2 cells in order to express relevant proteins that support the tissue during hair growth. Alternatively, SF1 and SF2 could be fuelled by two distinct cell populations which expand in either anagen or telogen skin. In-depth future studies are needed to investigate the function and lineage relationship of the identified fibroblast subtypes.

### A single-cell transcriptomic atlas of full-thickness skin during hair growth and rest

Finally, this comprehensive transcriptional atlas of full-thickness skin represents the diverse types of cells and their molecular programs in unprecedented detail. We made this unique resource freely and easily accessible through the web portal (http://kasperlab.org/tools) for further exploration to inform future research on skin and the stem and progenitor cell dynamics within the hair follicle.

## ACKNOWLEDGEMENTS

All single-cell experiments were performed with the help of the Eukaryotic Single Cell Genomics core facility at SciLifeLab, Stockholm. This work was supported by grants from the Swedish Research Council to MK and RS, the Swedish Cancer Society, Swedish Foundation for Strategic Research, Center for Innovative Medicine, and Ragnar Söderberg Foundation to MK, Karolinska Institutet PhD (KID) funding to SJ, TJ and KA, and a Wenner-Gren postdoctoral fellowship to TD. Parts of this study were performed at the Live Cell Imaging facility/ Nikon Center of Excellence, Department of Biosciences and Nutrition, Karolinska Institutet.

## AUTHOR CONTRIBUTIONS

MK and XS conceived the study. MK, XS, SJ, KA, TJ, TD and US planned the study details in general, and SJ envisioned and outlined all computational analyses. XS established the cell isolation protocol. XS, TJ and US performed cell isolation, and XS prepared cells for the 10 X Chromium system. KA performed smRNA-FISH staining with help from SJ. KA performed imaging and image preparation. SJ performed all data analysis with input/help from TJ (*Seurat* analyses). IS provided feedback throughout the entire course of the study. MK, RS, SJ, XS, IS, TD and KA interpreted the data. SJ prepared figures for tool. RS implemented tool online. SJ prepared all figures, tables and the first manuscript draft. MK and RS wrote the final manuscript version with input from SJ, KA, TJ, TD, XS and IS.

## METHODS

### Mouse work

All experiments were performed on female C57BL/6 mice. The mice were fed *ad libitum*, and handled and housed under standard conditions in the animal facility of Karolinska University Hospital Huddinge. All mouse experiments were performed in accordance to Swedish legislation and approved by the Stockholm South Animal Ethics Committee. Mice were sacrificed by cervical dislocation either during anagen at 5 weeks of age or during telogen at 9 weeks of age. Single-cell transcriptomes from n = 3 anagen and n = 2 telogen mice were included in the final dataset. Formalin-fixed paraffin-embedded (FFPE) sections were prepared from dorsal skin samples of the same mice.

### Cell isolation

After sacrifice, the back of the mice was shaved and the dorsal skin was isolated and floated on HBSS + 0.04% BSA. The skin was then cut into smaller pieces (around 1 mm in width) and incubated in 0.2% collagenase at 37°C for one hour. The partly digested skin was placed in a 70 µm cell strainer, smashed with a piston and rinsed three times with HBSS + 0.04% BSA. The flow-through was saved on ice, while the undigested tissue remaining in the cell strainer was incubated in 0.05% trypsin-EDTA for 15 minutes at 37°C. The trypsin-digested tissue was smashed once more and then passed through a 70 µm cell strainer. The flow-through of the collagenase digestion (containing mainly stromal cells) and the flow-through of the trypsin digestion (containing mainly epidermal cells) were pooled, spun down and subsequently filtered through a 40 µm cell strainer. The viability of the cell suspension was determined using trypan blue.

### Single-cell capturing, library preparation, generation and processing of sequencing data

Single-cell cDNA libraries were prepared using the Chromium Single Cell 3’ kit (version 1; 10X Genomics) according to the manufacturer’s instruction. The prepared libraries were sequenced on an Illumina HiSeq2500. Raw sequencing data were processed using the CellRanger package (v1.1.0; 10X Genomics). All downstream data analysis was performed using a mix of custom scripts and published analysis packages as described below.

### Data analysis

#### Analysis workflow

As described in **Figure S1C**, our analysis workflow encompassed two rounds of clustering and spatial embedding in combination with population-specific downstream analyses. For clustering and visualization, we used two approaches: (1) the default *Seurat* pipeline as gold standard in the field and (2) custom clustering with affinity propagation (AP) and visualization with UMAP. This tandem approach served as a qualitative control for cluster reproducibility and cluster robustness. 1^st^ level clustering served to define the main cellular compartments of the skin, while 2^nd^ level clustering was performed to identify compartment-specific subpopulations. As the aim of the 1^st^ level clustering was structuring the dataset in a biologically meaningful way, certain subpopulations already defined during the 1^st^ level clustering (e.g. different immune cell populations which were distinct enough to be separated in the first clustering round) were manually fused. 1^st^ and 2^nd^ level clustering was performed with both the *Seurat* and custom approach using the same cellular input. For the 1^st^ level clustering, the *Seurat* results are shown in **Figure 1** and served as input for the 2^nd^ level clustering, while the AP results are shown in **Figure S1E**. For the subclustering of permanent epidermis (EPI), anagen HF (ANA), fibroblasts and fibroblast-like cells (FIB), immune cells (IMM), vascular cells (VASC), neural-crested derived cells (NC) and miscellaneous cells (MISC), the AP results are shown in the main figures (**Figures 2-3, 5-6**) and served as input for the downstream analysis, while the Seurat results are shown in the respective supplementary figures (**Figures S2A, 5A, 6A, 7A-D**).

The following downstream analyses were performed:

(1) Test for statistical enrichment in anagen or telogen skin (all)
(2) Test for differential expression of genes in cell populations (all)
(3) Test for differential expression of genes between anagen and telogen (EPI)
(4) Prediction of cell cycle stage (all)
(5) Prediction of cell identity based on keratin expression (ANA)
(6) Analysis of RNA velocity with velocyto (ANA)
(7) Modeling gene expression along pseudotime (ANA)

All steps of the analysis workflow are described in detail below. Jupyter notebooks containing the full analysis workflow will be available upon publication.

#### Quality control

Overall, we observed that nearly all single-cell transcriptomes returned by the *CellRanger* package were of sufficient quality for downstream analysis. We for instance did not observe artificial cell clusters containing mostly low quality transcriptomes. Therefore, we did not perform any *a priori* thresholding based on for instance total number of molecules prior to clustering analysis. Instead we manually removed clusters containing biologically dubious signatures, most of which were generated by cell doublets, during the data analysis process.

#### Clustering and spatial embedding – Seurat pipeline

The *Seurat* package version 2.3.4 was used for 1^st^ and 2^nd^ level clustering. Gene expression data was preprocessed using log-normalization and regression of cell-to-cell variations driven by mitochondrial gene expression and the total number of detected molecules. For 1^st^ level analysis, linear dimensionality reduction was performed on the complete dataset using PCA with the most highly variable genes serving as input. The graph-based clustering approach encompassed building a KNN graph based on the Euclidian distance in PCA space, adjusting the edge weights using Jaccard similarity and clustering with the Louvain algorithm. t-SNE was used for visualization of the dataset. The same PCs were used as input for the clustering and the spatial embedding. During 2^nd^ level analysis, the same strategy for dimensionality reduction, clustering and visualization was applied. Variable genes and principle components were determined separately for each of the main cell types. Cluster resolution, t-SNE perplexity and the number of used principal components used for clustering and visualization were adjusted based on the total cell number of the respective cell types.

#### Clustering and spatial embedding – custom analysis pipeline

The custom analysis pipeline combines a variety of custom scripts and publically available analysis packages. Transcriptome normalization was based on the pool-based size factor approach described by (Lun, Bach, & Marioni, 2016a) and was performed using the *computeSumFactors* function of the *scran* R package (Lun, McCarthy, & Marioni, 2016b). High variance features were selected by fitting a second-degree polynomial to the variance as function of the mean. Non-linear dimensionality reduction was performed using Uniform Manifold Approximation (UMAP) implemented in *scanpy* (Wolf, Angerer, & Theis, 2018). Clustering was performed using *Affinity Propagation* (AP) as described before (Joost et al., 2016). Euclidean distance in PCA space served as input for both UMAP generation and AP clustering, with PCs chosen based on elbow plots. Cluster numbers were chosen based on Bayesian Information criterion (BIC) and biological considerations. Cells within clusters were ordered based on Ward linkage using POLO (Bar-Joseph, Gifford, & Jaakkola, 2001). To consolidate major discrepancies between AP clustering and UMAP embedding, i.e. individual or groups of cells removed from the majority of their AP cluster in the UMAP visualization, we used a reassignment approach based on k-nearest neighbors (kNN) in UMAP-space: a cell 𝑋 assigned to AP-cluster 𝐶𝑙(𝑋) was thereby reassigned if less than 𝑛 of its 𝑘-nearest neighbors (e.g. less than 15 of its 50 nearest neighbors) in UMAP-space belong to the same cluster 𝐶𝑙(𝑋), with the new cluster being based on the cluster assignment of the majority of its nearest neighbors.

#### Test of statistical enrichment in anagen or telogen skin

To test whether a certain cluster / population was enriched in anagen or telogen skin, we determined how likely the observed distribution of anagen and telogen cells could be the result of random sampling variation using a binomial distribution. To facilitate calculation using the cumulative density function, a population was considered enriched in telogen skin if:

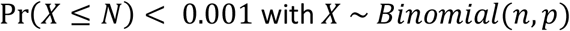

(i.e. it is unlikely that the same or fewer anagen cells are observed by chance).

Accordingly, a population was considered enriched in anagen skin if:

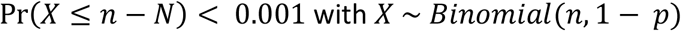

(i.e. it is unlikely that the same or fewer telogen cells are observed by chance).

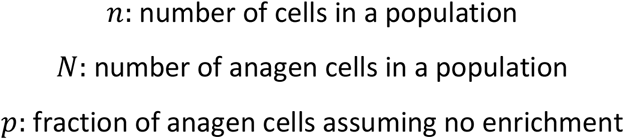

The value for 𝑝 - the expected fraction of anagen cells assuming no enrichment – is influenced by the fraction 𝑓_*a*_ and 𝑓*_t_* at which a cell type occurs in an anagen or telogen cell suspension over replicates 𝑟 = 1 − 𝑖, and the number of sequenced cells 𝑆*_a_* and 𝑆*_t_* over replicates 𝑟 = 1 − 𝑖. Thus

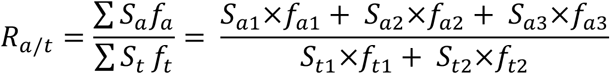

Under the null hypothesis that a cell type is present in equal total numbers (per area) in anagen and telogen skin, 𝑓*_a_* and 𝑓_*t*_ an be defined as 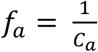 and 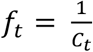 respectively, where 𝐶*_a_* and 𝐶*_t_* are the total number of cells isolated from an area of skin assuming that experimental conditions are similar. While it is possible to use values for 𝐶*_a_* and 𝐶*_t_* for each replicate 𝑟 = 1 – 𝑖 individually, we assume that differences between replicate values for 𝐶 represent experimental noise and do not reflect true differences of 𝑓 in the tissue. We thus use mean values for 𝐶*_a_* and 𝐶*_t_* over all replicates to approximate true values for 𝑓*_a_* and 𝑓*_t_*. Thus

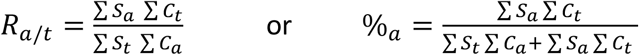

As we observed average cell numbers of 𝐶*_t_* = 4.*7* × 10^5^ and 𝐶*_a_* = 13.*2* × 10^5^ in telogen and anagen respectively, and sequenced 2764 telogen and 3331 anagen cells, we assume that 𝑝 = %*_a_* = 0.30. Reassuringly, the median values for %_*a*_ among 1^st^ and 2^nd^ level clusters are very similar to this inferred value (**Figure S1D**).

#### Test for differential expression of genes in cell populations

Genes significantly overexpressed in a cell population were identified with the Mann-Whitney U / Wilcoxon rank sum test using the *scipy* implementation. Gene expression in a cell population was either compared to a pool of all other populations, or to each other population individually with the largest P-value returned (i.e. in Figure S1J the differential expression compared to the most similar main cluster is plotted). Correction for multiple testing was performed using the Benjamini-Hochberg method. The significance threshold was set to 𝛼 = 0.001 unless specified differently.

#### Test for differential expression of genes between anagen and telogen

Due to the overall lower number of unique molecules detected in telogen cells compared to anagen cells (compare **Figure S1B**), tests for differential expression between anagen and telogen are susceptible to bias even after normalization due to the higher degree of zero inflation in telogen cells. We thus downsampled all permanent epidermis cells to 1000 counts (using the *scanpy* implementation) before performing differential expression analysis between anagen and telogen as described above. The significance threshold was set to 𝛼 = 0.05.

#### Prediction of cell cycle stage

Cell cycle stage was predicted using the *CellCycleScoring* function implemented in the *Seurat* package (Butler et al., 2018).

#### Prediction of cell identity based on keratin expression

In order to predict anagen HF layer identity based on keratin expression, layer specific-expression patterns were taken from (Langbein et al. 2010). Anagen HF single-cell transcriptomes were then scored for layer identity based on their keratin expression signatures using an approach similar to *RaceID* (Grün et al., 2015) and the Naïve Bayes method introduced in (Joost et al., 2018). In brief, a second-degree polynomial was fitted to the relationship between mean expression and variance considering all genes expressed in anagen HF cells. Based on this fit, mean and expected variance for each keratin gene 𝑔 were determined and used to fit a negative binomial representing background expression

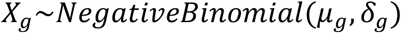

For each anagen HF transcriptome 𝑋*_i_*, the probability Pr(𝑋*_g_* ≥ 𝑋*_ig_*) of an expression value of 𝑋*_ig_* or more extreme being generated from the background distribution was determined. Layer-specific scores 𝑆*_i_* rewarding above-background expression of layer-specific keratins and penalizing above-background expression of keratins not expressed in a layer were then calculated as

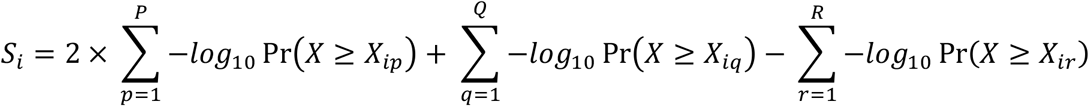

where 𝑃 contains all keratin genes uniquely expressed in a layer (given twice the weight), 𝑄 contains all keratin genes non-uniquely expressed in a layer and 𝑅 contains all keratin genes not expressed in a layer.

#### Analysis of RNA velocity with velocyto

To establish whether a differentiation relationship is present in anagen HF cells, we estimated RNA velocity using the *velocyto* package (La Manno et al., 2018). Counts of unspliced and spliced mRNA in anagen cells were generated using the *velocyto* CLI according to standard protocol. RNA velocity was subsequently estimated in all anagen HF cells (‘all), all ORS cells (‘ORS’) or all *Msx2*+ anagen HF cells (‘matrix’) using the same workflow and similar parameters. In brief, normalization by total expression was performed after selecting genes (*min_expr_counts = 10*, *min_expr_counts_U = 1*, *min_cells_express_U = 1*), kNN imputation was then performed in PCA space (‘all’: k = 100, n_components = 15; ‘ORS’: k = 25, n_components = 5, ‘matrix’: k = 100, n_components = 15) and gammas were fitted using *weights = ‘maxmin_diag’*. Genes were filtered based on *minR2 ≥ 0.2* and *min_gamma ≥ 0.01*. RNA velocity was subsequently estimated assuming constant velocity and transition probability as well as embedding shift were calculated based on the previously generated UMAP representation of the anagen dataset (transform = “log”, n_sight = 500 (‘all’,’matrix’) or 100 (‘ORS’), sigma_corr = 0.1). The Markov models were created using similar steps as presented in the (La Manno et al., 2018) paper.

#### Modeling gene expression along pseudotime

Diffusion maps and diffusion pseudotimes for *Msx2*+ anagen HF cells and anagen HF IRS cells were created using the *scanpy* implementation (Haghverdi et al., 2016; Haghverdi, Buettner, & Theis, 2015; Wolf et al., 2018). For *Msx2*+ anagen HF and anagen HF IRS cells, outlier cells were removed by creating a minimum spanning tree in diffusion space, defining diameter paths through all branches and eliminating cells distant to the diameter paths. Root cells for the estimation of diffusion pseudotime were then defined based on the *velocyto* Markov models (cells with maximum density in ‘backwards’ models) presented in **Figure 3D**. Diffusion pseudotime branches were separated manually based on clustering data projected on diffusion maps.

Gene expression along pseudotime in *Msx2*+ anagen HF cells and anagen HF IRS was modeled as a generalized linear model (GLM) using the R package *VGAM* in *rpy2* based on an approach first introduced by (Trapnell et al., 2014) and (Qiu et al., 2017). For each gene, gene expression was modeled along pseudotime as a natural spline with 3 degrees of freedom (full model) and compared to several reduced and null models:

##### Full model assuming branch-specific, pseudotime-dependent expression

𝑌 ∼ 𝑛𝑠 𝑋, 𝑑𝑓 = 3 × 𝐵, where 𝑋 is the position of cells along pseudotime, 𝑌 the expression of a gene and 𝐵 a branch-specific binary predictor.

##### Reduced model assuming branch-independent, pseudotime-dependent expression

𝑌 ∼ 𝑛𝑠 𝑋, 𝑑𝑓 = 3, where 𝑋 is the position of cells along pseudotime and 𝑌 the expression of a gene.

##### Null model assuming constant expression at level of root cell population

𝑌 ∼ 𝐵, where 𝑌 is the expression of a gene and 𝐵 a binary predictor specifying cells belonging to the root cell population (i.e. the AP cluster that the root cell is assigned to).

To establish pseudotime-dependency, the full model was compared to the null model and to establish branch-specificity, the full model was compared to the reduced model using the likelihood ratio test implemented in *VGAM*. For a gene being considered as differentially regulated over pseudotime, the following conditions must have been met:

##### *Msx2*+ anagen HF cells

(1) *max expression (fitted) >* 0.25
(2) *likelihood ratio test (full, null)* < 0.001
(3) *fold change between max and min expression >* 2
(4) *fold change between max expression and null mode*l > 2

##### Anagen HF IRS cells

(1) *max expression (fitted) >* 0.25
(2) *likelihood ratio test (full, null)* < 0.01
(3) *Fold change (fc) between max and min expression >* 2
(4) *fc between max expression and null mode*l > 2

Pseudotime-dependent genes were clustered into gene modules based on Pearson correlation of fitted natural splines using *Affinity Propagation*.

### Fluorescent *in situ* hybridization (FISH)

RNA-FISH was performed using the RNAscope® Multiplex Fluorescent Detection Kit v2 (323100, Advanced Cell Diagnostics) according to manufacturer’s instructions using TSA with Cy3, Cy5, and/or Fluorescein (NEL760001KT, Perkin Elmer) on FFPE sections of anagen and telogen skin. All sections were counterstained with either WGA (W11261, Invitrogen), DAPI (D1306, Invitrogen) or both. Each individual staining and/or co-staining was performed on skin samples from at least 3 different mice for anagen skin (Figures 2, 3, 4, 5, S2, S5 and S6), and at least 2 different mice for telogen skin (Figures 5 and S5). Images were acquired on a Nikon A1R spinning disk confocal or Zeiss LSM710 confocal and were all processed the same way (maximum intensity projection, background removal with the “subtract background” plug-in, brightness adjustment, pseudocoloring) using Fiji (Rueden et al., 2017; Schindelin et al., 2012). The following probes were used:

| Probe | Product number |
| --- | --- |
| *Barx2* | 548011-C3 |
| *Cd34* | 319161-C2 |
| *Cited4* | 553891 |
| *Dcn* | 413281 |
| *Edn2* | 418221 |
| *Fgf5* | 417091 |
| *Foxp1* | 485221-C2 |
| *Krt6a* | 464441 |
| *Krt75* | 430121-C2 |
| *Krt79* | 436201-C2 |
| *Lef1* | 441861 |
| *Lgr5* | 312171-C2 |
| *Mfap5* | 490211-C2 |
| *Msx2* | 421851 |
| *Foxq1* | 504801 |
| *Gata3* | 403321-C3 |
| *Gpx3* | 509901-C3 |
| *Id3* | 445881-C2 |
| *Il11ra1* | 548261-C2 |
| *Krt5* | 415041-C2 |
| *PanHh* | 415051-C2 |
| *Plac8* | 532701 |
| *Pou3f1* | 436421-C2 |
| *Pthlh* | 456521-C2 |
| *Shh* | 314361-C2 |
| *Sparc* | 466781-C2 |

## DATA AND SOFRWARE AVAILABILITY

### Data resources

The sequencing data has been deposited at the Gene Expression Omnibus (GEO) with the accession number GSE129218, and will be available upon publication.

### Software

The complete computational analysis workflow will be available in the form of jupyter notebooks and deposited upon publication.

---

**Figure S1.**
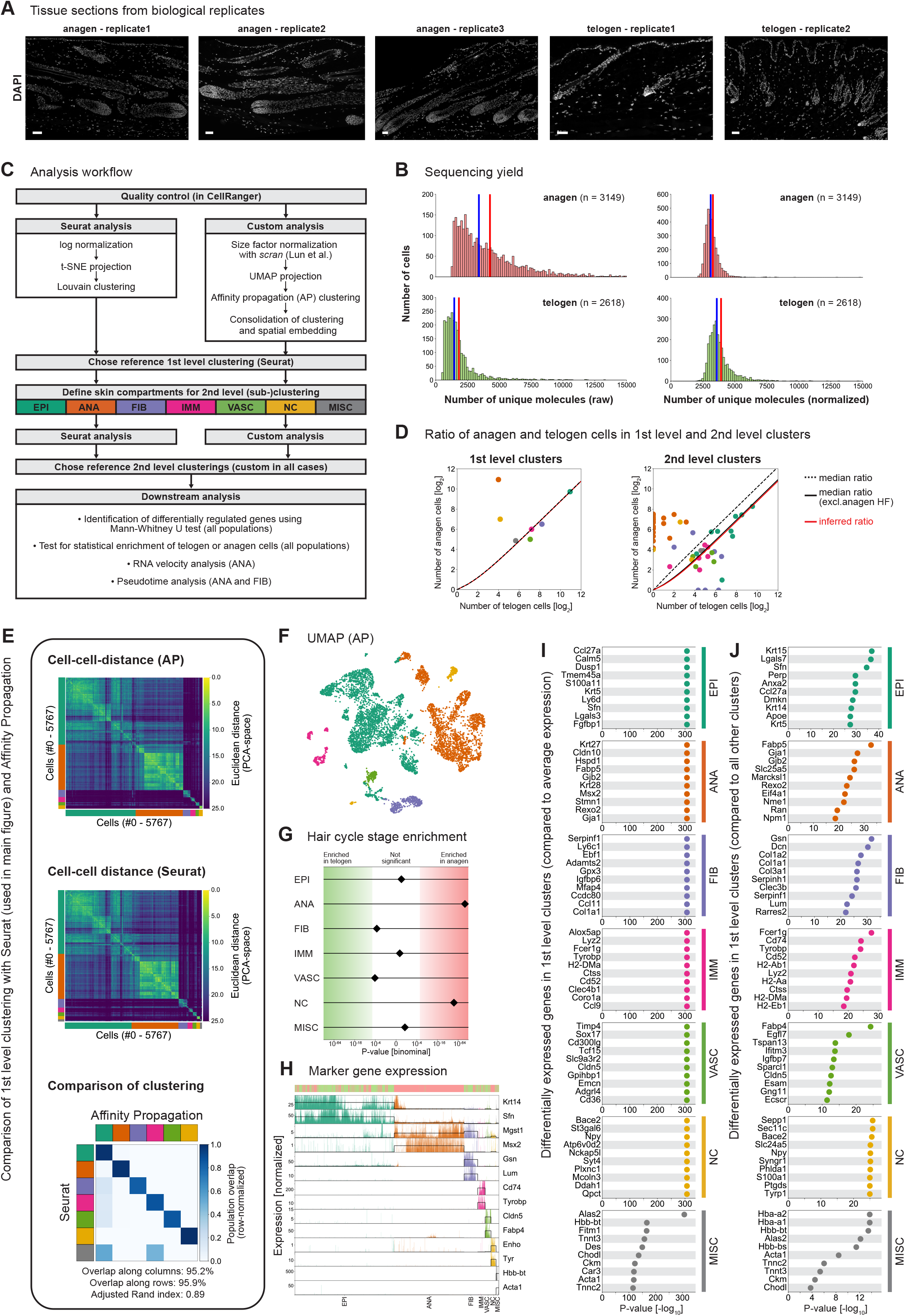
Details on the clustering of single-cell RNA-seq data from anagen and telogen skin. Related to Figure 1. (**A**) DAPI stained tissue sections from each biological replicate showing hair cycle stage. (**B**) Histograms showing the number of unique molecules detected in anagen or telogen data before (left) and after size factor normalization (right). Red and blue lines denote mean and median respectively. (**C**) Flow diagram summarizing the analysis workflow. (**D**) Ratio of anagen and telogen cells in 1^st^ and 2^nd^ level clusters. Red lines show the ratio of anagen and telogen cells expected to be observed in a certain population in the data if the populations would be of equal size (cells / area) in anagen and telogen skin (compare Methods). Dotted black lines show the median anagen / telogen ratio observed in the data, while the filled black line shows the median anagen / telogen ratio observed in the 2^nd^ level data if anagen HF populations (highly anagen enriched) are not considered. (**E**) Comparison of 1^st^ level clustering using affinity propagation (AP) with the *Seurat* approach shown in Figure 1 (visualization with t-SNE; clustering using Louvain). Upper panel: Euclidean distances (considering 30 PCs) of cells ordered according to AP clustering. Centre panel: Euclidean distances (considering 30 PCs) of cells organized according to *Seurat* clustering (with in-cluster order defined by Ward linkage clustering). Lower panel: Comparison of *Seurat* clustering with AP-clustering. Shown is the row-normalized overlap between clusters. Maximal overlap was calculated row-wise and column-wise as the optimal overlap (shared number of cells) of each *Seurat* (row) and AP cluster (columns) divided by the total number of cells. (**F**) AP clusters projected onto UMAP visualization of whole dataset. (**G**) Statistical test for enrichment of main populations (1^st^ level clusters) in either anagen or telogen skin. Shown is the probability (P-value) that the anagen/telogen composition of a population can be observed by chance assuming no enrichment (but controlling for the expansion of anagen skin). If the P-value exceeds the threshold of 𝛼 = 0.001 (Bonferroni-corrected), a population is assumed to be statistically significantly enriched in either anagen or telogen skin. (**H**) Expression of marker genes associated with each main population. Each bar represents a single-cell transcriptome, line denotes median expression within clusters. Hair cycle stage of cells is shown in the top panel (green: telogen; red: anagen). (**I, J**) Most significantly overexpressed genes (P-value < 0.001) in each main population (I) compared to average expression over all other populations or (J) compared to each other population separately (highest P-value is returned). Note that in order to avoid infinite values in negative logarithmic representation, P-values = 0 have been set to 1e-307. Scale bars: 50 µm (A).

**Figure S2.**
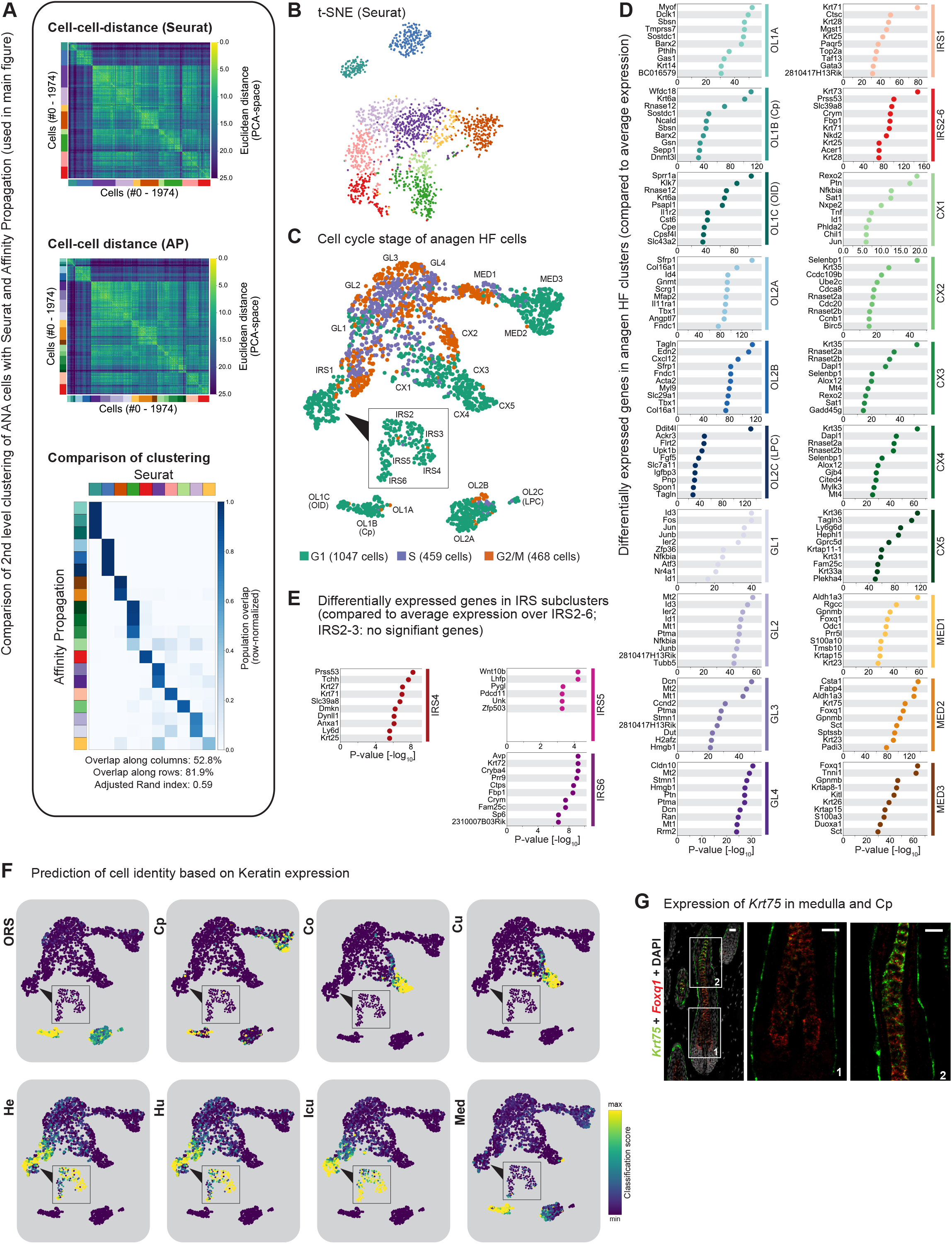
Subclustering of anagen HF cells. Related to Figures 2 and 3. (**A**) Comparison of anagen HF subclustering using *Seurat* with the AP approach shown in Figures 2 and 3. Upper panel: Euclidean distances (considering 20 PCs) of cells ordered according to *Seurat* clustering. Centre panel: Euclidean distances (considering 20 PCs) of cells ordered according to AP clustering. Lower panel: Comparison of *Seurat* clustering with AP clustering. Shown is the row-normalized overlap between clusters. Maximal overlap was calculated row-wise and column-wise as the optimal overlap (shared number of cells) of each AP (row) or *Seurat* cluster (columns) divided by the total number of cells. (**B**) *Seurat* clusters projected onto t-SNE visualization of anagen HF. (**C**) Predicted cell cycle stage projected onto UMAP representation of anagen HF cells introduced in Figure 2A. (**D-E**) Most significantly overexpressed genes (P-value < 0.001) in each (D) anagen HF subpopulation or (E) IRS subpopulation compared to average expression over all other populations (within the ANA main cluster). (**F**) Classification of cell identity based on keratin expression (taken from (Langbein et al., 2010)) observed in different compartments of the anagen HF. (**G**) smRNA-FISH staining of Cp marker *Krt75* and MED marker *Foxq1* (n = 3 mice). Scale bars: 20 µm.

**Figure S3.**
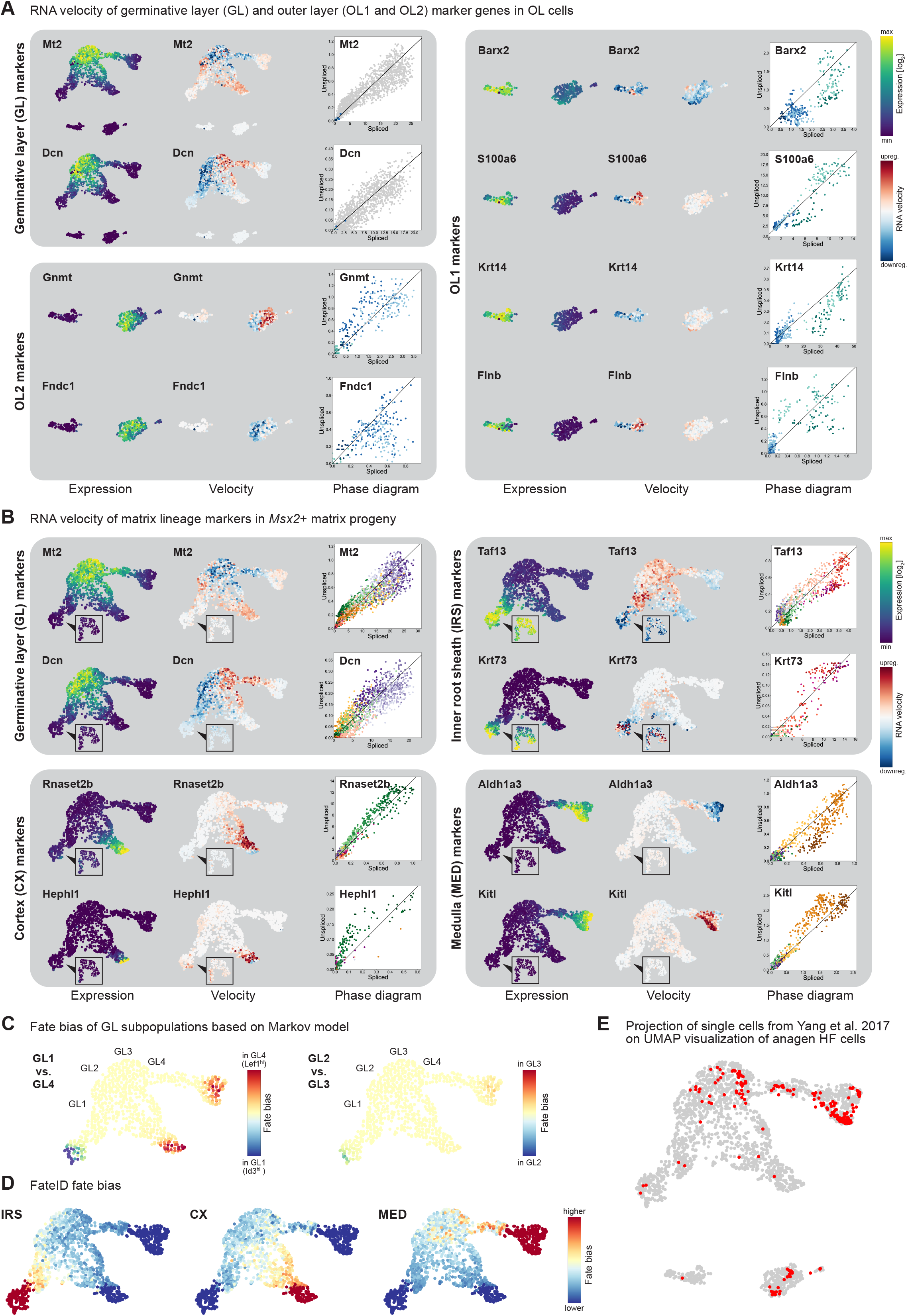
Analysis of matrix progeny differentiation with *velocyto*. Related to Figures 2 and 3. (**A-B**) RNA velocity analysis of selected genes linked to the OL1, OL2 or GL compartment (A) or the matrix progeny compartments (B) of the anagen HF. Shown for each gene is (left) the kNN-imputed expression of spliced mRNA, (centre) the cell-specific RNA velocity inferred from its unspliced-spliced ratio and (right) the phase diagram showing the unspliced-spliced ratio over all included cells with the fitted line denoting the steady state. In the phase diagram cells are coloured according to cluster identity (as in Figures 2E and 3A). To promote visibility of OL cells (in A), *Msx2*+ cells have been greyed out in the phase diagrams of GL associated genes. UMAPs are the same as introduced in Figure 2A, 2E or 3A. (**C**) Fate of germinative layer cells (GL1-GL4) based on Markov model (forward) of RNA velocity data. Shown are differences in cell fate originating from different GL subpopulations. (**D**) Bias towards IRS, CX or MED lineage determined by FateID (Herman et al., 2018). (**E**) Projection of anagen HF cells from (Yang et al., 2017) (red) onto UMAP introduced in Figure 2A.

**Figure S4.**
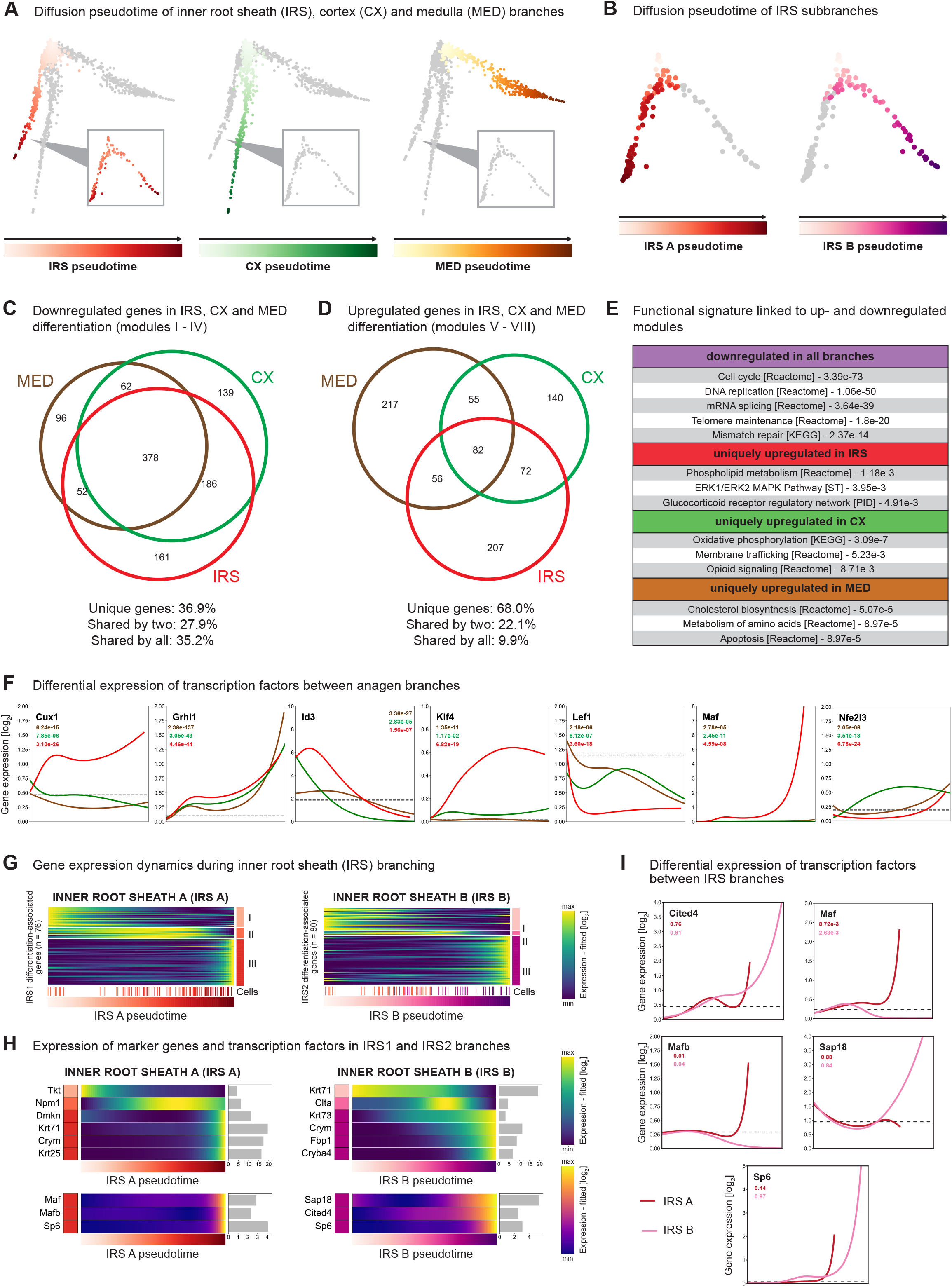
Pseudotime modelling of matrix differentiation. Related to Figure 4. (**A**) Diffusion pseudotime of IRS, CX and MED branches projected onto diffusion map introduced in Figure 4A. The root cell required for the generation of diffusion pseudotime was chosen based on the Markov model in Figure 3E (cluster: GL4). (**B**) Diffusion pseudotime of IRS subbranches projected onto diffusion map introduced in Figure 4A. Root cell was chosen based on Markov model in Figure 3D (cluster: IRS2). (**C**) Venn diagram comparing genes downregulated in IRS (red), CX (green) and MED (brown) branches (Modules I – IV). (**D**) Venn diagram comparing genes transiently or permanently upregulated in IRS (red), CX (green) and MED (brown) branches (Modules V-VIII). (**E**) Functional signatures linked to genes downregulated in all branches, or genes upregulated uniquely in IRS, CX, or MED branches. (**F**) Comparative expression (fitted) of transcription factors in IRS (red), CX (green) and MED (brown) branches. Dotted lines show the null model (baseline expression in root cell cluster GL4). Numbers denote the P-values for branch-specificity compared to a reduced model which assumes equal expression over all branches. Note: A low P-value can signify both increased and decreased branch-specific expression compared to the reduced model. (**G**) Gene expression dynamics along IRS subbranches. Shown in the heatmap are the expression patterns (fitted) of significantly pseudotime-dependent genes divided into three gene modules. The lower panels show the location of individual cells along pseudotime. (**H**) Expression patterns (fitted) of selected marker genes and transcription factors along pseudotime. Shown in the right panels are the P-values for pseudotime dependency compared to a null model assuming no change in expression compared to the root cell cluster (GL4). (**I**) Comparative expression (fitted) of transcription factors in IRS A (red) and IRS B (pink) subbranches. Dotted lines show the null model (baseline expression in root cell cluster IRS2). Numbers denote the P-values for branch-specificity compared to a reduced model which assumes equal expression over all branches. Note: A low P-value can signify both increased and decreased branch-specific expression compared to the reduced model.

**Figure S5.**
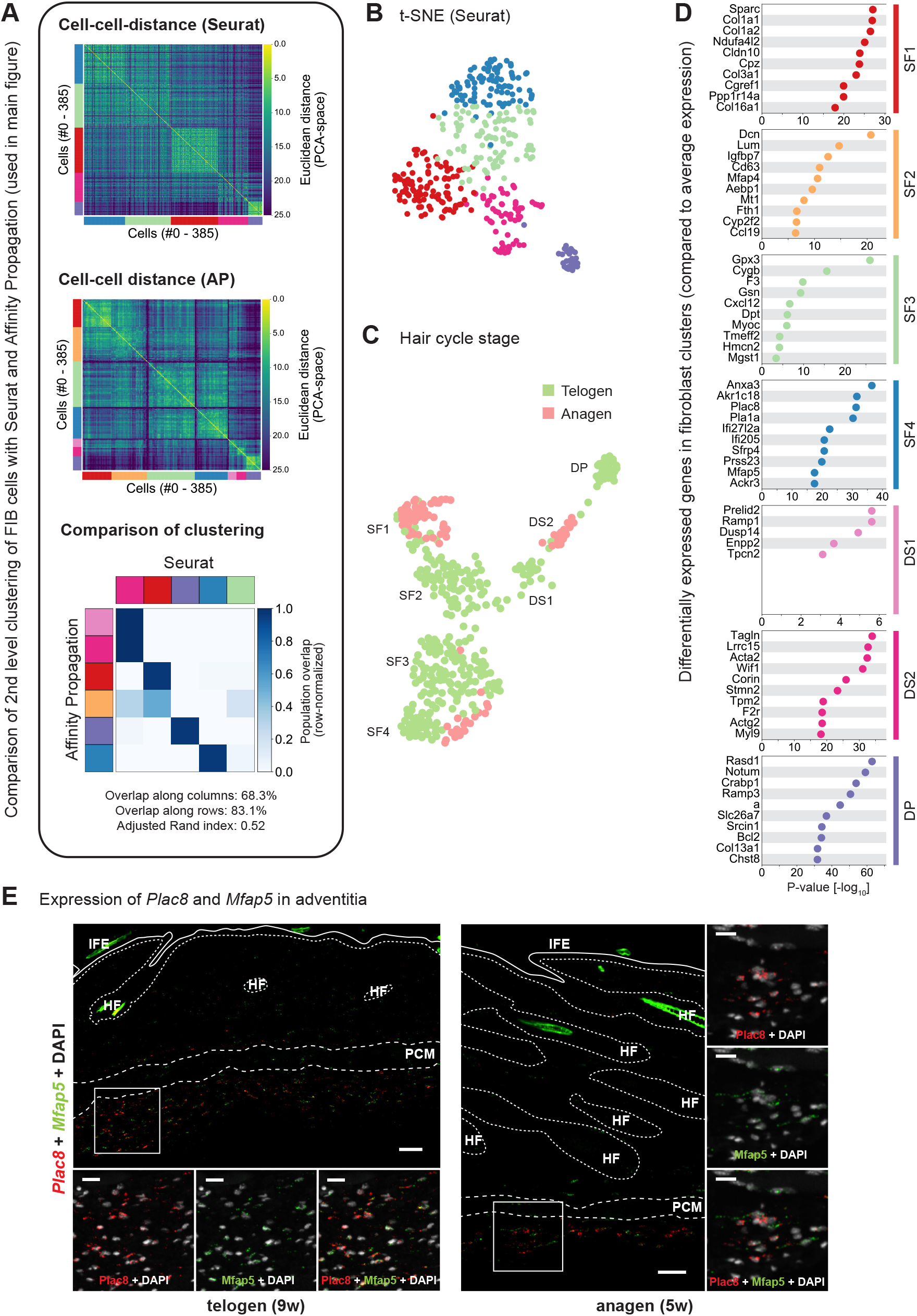
Subclustering of fibroblasts and fibroblast-like cells. Related to Figure 5. (**A**) Comparison of fibroblast subclustering using *Seurat* with the AP approach shown in Figure 5. Upper panel: Euclidean distances (considering 15 PCs) of cells ordered according to *Seurat* clustering. Centre panel: Euclidean distances (considering 15 PCs) of cells ordered according to AP clustering. Lower panel: Comparison of *Seurat* clustering with AP clustering. Shown is the row-normalized overlap between clusters. Maximal overlap was calculated row-wise and column-wise as the optimal overlap (shared number of cells) of each AP (row) or *Seurat* cluster (columns) divided by the total number of cells. (**B**) *Seurat* clusters projected onto t-SNE visualization of fibroblasts. (**C**) Hair cycle stage projected onto UMAP representation of fibroblasts introduced in Figure 5A. (**D**) Most significantly overexpressed genes (P-value < 0.001) in each fibroblast population compared to average expression over all other populations (within main population). (**E**) smRNA-FISH staining of SF4 markers *Plac8* and *Mfap5*. Shown are overviews without and zoom-ins with nuclear stain (n = 3 mice for anagen, n = 2 mice for telogen). HF: Hair follicle. IFE: Interfollicular epidermis. PCM: *Panniculus carnosus* muscle. Scale bars: 50 μm (overviews), 20 µm (insets) (E).

**Figure S6.**
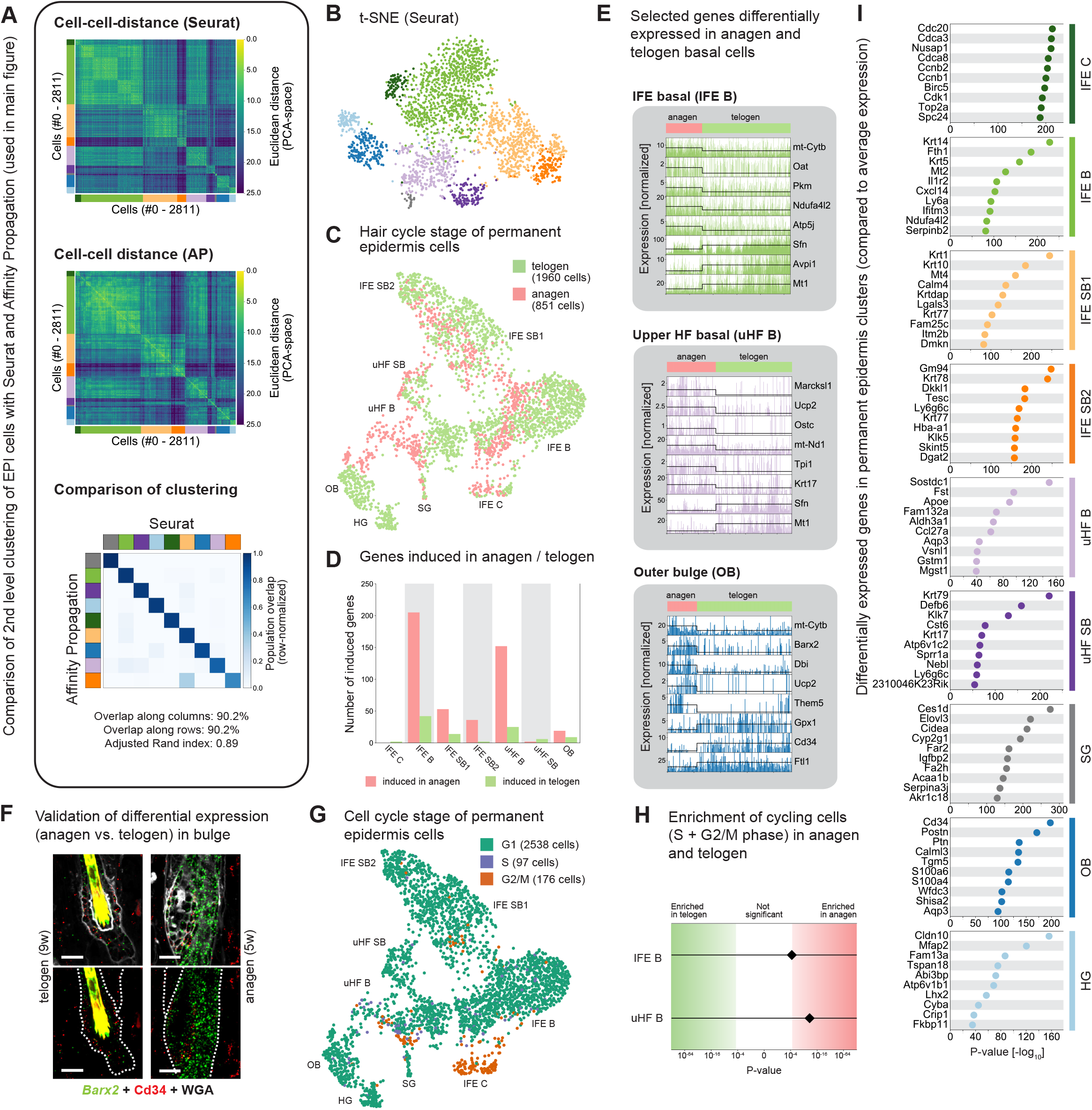
Subclustering of permanent epidermis cells. Related to Figure 6. (**A**) Comparison of permanent epidermis subclustering using *Seurat* with the AP data shown in Figure 6A. Upper panel: Euclidean distances (considering 15 PCs) of cells ordered according to *Seurat* clustering. Centre panel: Euclidean distances (considering 15 PCs) of cells ordered according to AP clustering. Lower panel: Comparison of *Seurat* clustering with AP clustering. Shown is the row-normalized overlap between clusters. Maximal overlap was calculated row-wise and column-wise as the optimal overlap (shared number of cells) of each AP (row) or *Seurat* cluster (columns) divided by the total number of cells. (**B**) *Seurat* clusters projected onto t-SNE visualization of epidermal cells. (**C**) Hair cycle stage projected onto UMAP representation of permanent epidermis introduced in Figure 6A. (**D**) Number of genes significantly overexpressed (P-value < 0.05) in anagen or telogen cells of epidermal populations based on a downsampling model. (**E**) Expression of selected genes that are significantly differentially regulated between anagen and telogen in IFE basal (upper), upper HF (centre) and outer bulge cells (lower). Each bar represents a single-cell transcriptome, line denotes median expression within clusters. While expression levels after size factor normalization are shown, the presented genes were identified using a downsampling approach as described in the Methods. (**F**) smRNA-FISH staining of *Barx2* and *Cd34* in telogen and anagen bulges (n = 3 mice for anagen, n = 2 mice for telogen). Scale bar: 20 µm. (**G**) Cell cycle stage classification projected onto UMAP introduced in Figure 6A. (**H**) Statistical test for enrichment of proliferative IFE basal and upper HF cells in anagen or telogen skin. Cells with either an S or G2/M classification as shown in Figure S6C were considered proliferative. (**I**) Most significantly overexpressed genes (P-value < 0.001) in each epidermal population compared to average expression over all other populations (within main population).

**Figure S7.**
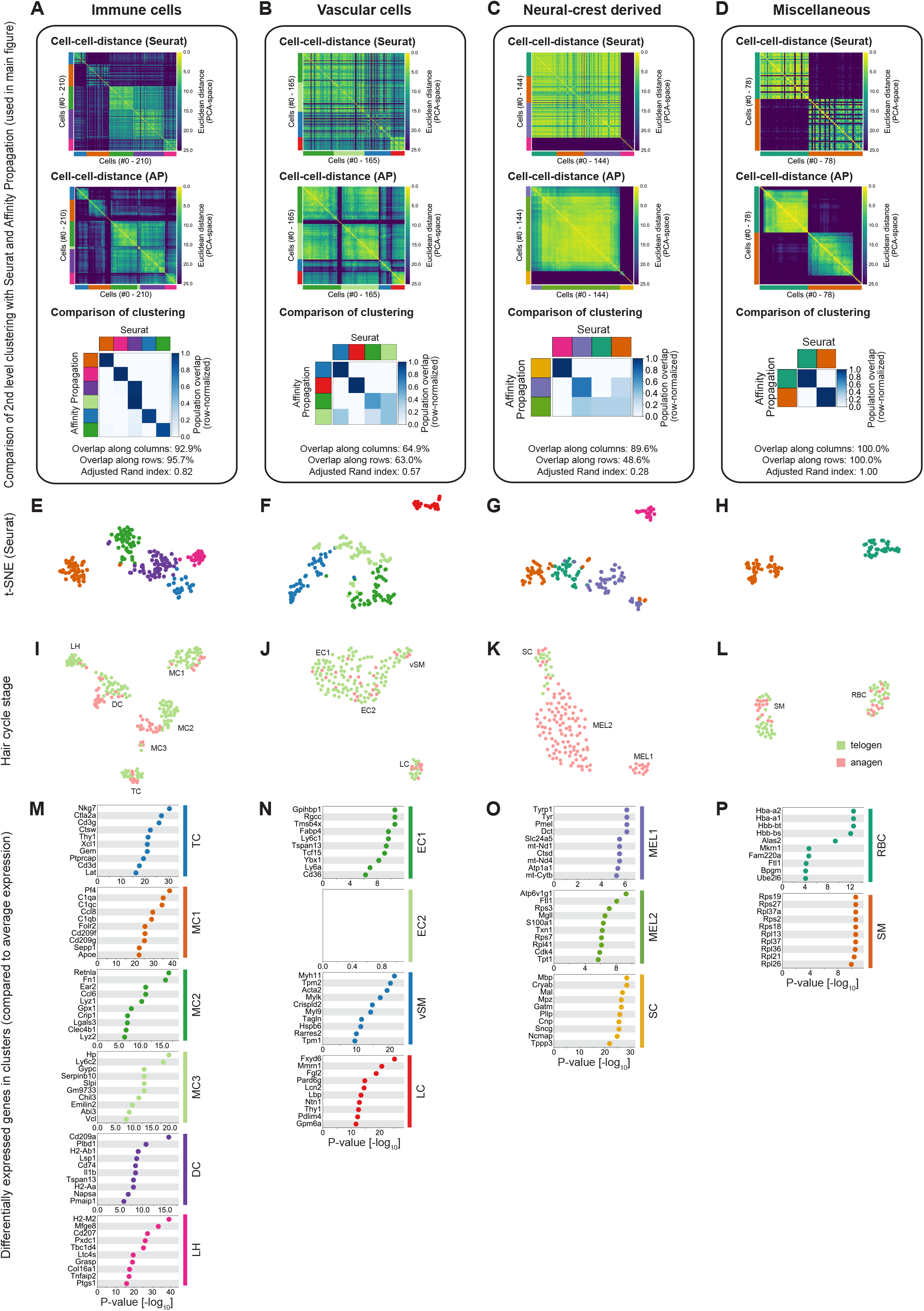
Subclustering of remaining stromal compartments. Related to Figure 6. (**A-D**) Comparison of immune cell (A), vascular cell (B), neural crest-derived cell (C) and miscellaneous cell (D) subclustering using *Seurat* with the AP approaches shown in Figure 6. Upper panel: Euclidean distances (considering 15 PCs, 8 PCs, 15 PCs and 10 PCs respectively) of cells ordered according to *Seurat* clustering. Centre panel: Euclidean distances (considering 15 PCs, 8 PCs, 15 PCs and 10 PCs respectively) of immune cells, vascular cells, neural crest-derived cells and miscellaneous cells. Lower panel: Comparison of *Seurat* clustering with AP clustering. Shown is the row-normalized overlap between clusters. Maximal overlap was calculated row-wise and column-wise as the optimal overlap (shared number of cells) of each AP (row) or *Seurat* cluster (columns) divided by the total number of cells. (**E-H**) *Seurat* clusters projected onto t-SNE visualization of immune cells (E), vascular cells (F), neural crest-derived cells (G) and miscellaneous cells (H) ordered according to AP clustering. (**I-L**) Hair cycle stage projected onto UMAP representation of immune cells (I), vascular cells (J), neural crest-derived cells (K) and miscellaneous cells (L) introduced in Figure 6B-D. (**M-P**) Most significantly overexpressed genes (P-value < 0.001) in each immune cell (M), vascular cell (N), neural crest-derived cell (O) and miscellaneous cell (P) population compared to average expression over all other populations (within respective main population).

